# Population-based body-brain mapping links brain morphology and body composition

**DOI:** 10.1101/2020.02.29.970095

**Authors:** Tiril P Gurholt, Tobias Kaufmann, Oleksandr Frei, Dag Alnæs, Unn K Haukvik, Dennis van der Meer, Torgeir Moberget, Kevin S O’Connell, Olof D Leinhard, Jennifer Linge, Rozalyn Simon, Olav B Smeland, Ida E Sønderby, Adriano Winterton, Nils Eiel Steen, Lars T Westlye, Ole A Andreassen

## Abstract

**Background:** Understanding complex body-brain processes, and putative interplay between adipose tissue and brain health, is of vital importance for brain and somatic disease prevention in the general population. We studied the link between body composition and brain structure through large-scale investigation in a healthy population without secondary disease effects.

**Methods:** We processed brain magnetic resonance imaging (MRI) data and extracted measures of brain morphometry from 19,330 healthy UK Biobank participants, of which a subset (n=2,703) had body MRI. We investigated associations between brain structure and (i) anthropometric body composition measures, and (ii) regional/specific body MRI measures of abdominal fat and muscle tissue.

**Findings:** We identified highly significant body-brain associations (*p*-values≤0·0002). Anthropometric measures showed negative, nonlinear, associations with cerebellar/cortical gray matter, and brain stem structures, negative associations with white matter, and positive associations with ventricular volumes. Subcortical structures exhibited mixed effect directionality, with strongest positive association for accumbens. Among body MRI measures, liver fat was negatively associated with thinner/lower cortical gray matter thickness/volume, and thigh muscle volume positively associated with accumbens volume.

**Interpretation:** We demonstrate significant body-brain associations, and map individual differences in body composition to brain morphology in healthy individuals. Common measures of body composition correlated negatively with cerebellar and cortical structures and positively with the accumbens, a dopamine rich structure involved in reward processing. These findings of a relationship between brain anatomy and body composition provide new insight into body-brain processes and suggest shared mechanisms of cardiometabolic risk factors and brain disorders. This may form the foundation for a new type of prevention studies, and provides a framework for studies of underlying mechanisms related to unhealthy lifestyle and obesity, with implications for public health and prevention.

**Funding:** The Research Council of Norway, South-Eastern Norway Regional Health Authority, European Union’s Horizon 2020 Research and Innovation Programme & European Research Council.

**Research in context:** *Evidence before this study:* Prior studies have indicated an association between brain structure and both obesity and fitness levels - of opposing directionality. Despite this, normal body-brain association patterns in healthy individuals have not been established, and the causal mechanisms are unclear. To enhance our understanding and establish the link between the body and the brain, we saw the need for large-scale investigations in healthy populations. For the study, we searched the PubMed database from March 12^th^, 2019, through February 25^th^, 2020, for scientific literature related to adipose tissue, body composition, brain morphology, and body and brain MRI. Search terms included: body fat, adipose tissue, subcutaneous/visceral adipose tissue, liver fat, body composition, anthropometric measures, body mass index, waist circumference, waist-to-hip ratio, adiposity, obesity, metabolic syndrome, cardiovascular, cardiometabolic, disease/disorder, muscle volume, fitness, brain structure, brain morphology, brain MRI, and body MRI. We based the scientific foundation on review studies, meta-analyses, and other larger studies, but generally excluded smaller studies, and thereby lowering the risk of evidential bias such as winner’s curse, although this does not eliminate the risk of publication bias.

*Added value of this study:* In the largest study, to date, including 19,330 healthy participants without secondary disease effects, we provide insight into normal body-brain processes by identifying body-brain associations that map normally varying body composition to brain morphology.

*Implications of all the available evidence:* We identified body-brain associations that give insight into normal physiological body-brain processes in healthy individuals, providing a reference point for studies of underlying mechanisms related to unhealthy lifestyle, obesity, and disorders of the body and the brain. Whereas the directionality and causal chain is unknown, these findings have potential implications for public health and disease prevention.

## Introduction

Obesity is a risk factor for disorders of both the body^1^ and the brain,^1,2^ and represents a global health challenge. Although causal mechanisms remain unclear, the associations between brain and physical health likely reflect body-brain interactions and common mechanisms across the soma and the mind. Indeed, patients with mental disorders show subtle structural brain alterations as revealed using brain imaging,^3^ and are at increased risk for poor physical health, including obesity, metabolic syndrome, cardiovascular disorders, and shorter life-expectancy.^4^ Yet, how these factors relate to brain health remains poorly understood.

Cross-sectional brain magnetic resonance imaging (MRI) studies have documented negative associations between obesity/poor metabolic health and gray matter volumes^5–9^ and white matter microstructure,^5,6^ while there is conflicting evidence for white matter volume.^5,6^ It is not clear how different aspects of obesity, e.g. adipose tissue distribution, or intra-abdominal fat - a known risk factor for adverse health outcomes^1,10^ - relate to brain health. Nonlinear associations are a common phenomenon in neuroimaging (e.g. accelerated brain atrophy at higher ages^11^), yet it is unknown whether aspects of obesity are nonlinearly related to brain structure. The genetic contribution to obesity is substantial and interacts with the environment, lifestyle, and sex.^12^ Additionally, body composition/obesity, as well as brain structure, are modifiable and sensitive to environmental and lifestyle factors, and sex. Indeed, physical fitness and activity have been positively associated with gray and white matter volumes^13^, indicating their importance for brain health, while negative associations have been reported for mobility impairment.^14^

Prior studies have largely investigated associations between brain structure and anthropometric measures (e.g. body mass index (BMI), waist-to-hip ratio (WHR)), which are informative, but nonspecific measures of body composition. In contrast, body MRI enables more specific and detailed *in vivo* measures of regional fat and muscle distribution, allowing for individual body composition profiling which has relevance for clinical prediction.^15^ How, and to what degree, these body MRI measures relate to individual differences in brain structure is unknown.

The pathophysiological mechanisms of psychiatric disorders, and other brain disorders, have proven elusive. To disentangle these complex and multifactorial mechanisms, with each contributing factor having small effects, we need an improved foundational understanding of normal body-brain connections in healthy individuals. To accurately capture such small effects, and thereby improve our understanding of normative body-brain connections, large-scale investigations are needed.^16^

To this end, we tested for associations between body composition and brain structure in healthy individuals aged 44 to 82 years using anthropometric measures and brain (n=19,330) and body (n=2,703) MRI from the UK biobank.^17^ Based on prior studies^5–9,13^ we expected brain structure to show negative associations with body composition measures, possibly with stronger associations for intra-abdominal fat, and positive associations with muscle volume. Through large-scale mapping, we hypothesized that we would obtain insight into, and identify robust patterns of normal body-brain connections, in a healthy population.

## Methods

### Study design and participants

We included 19,330 healthy UK biobank participants (10,104 women, 9,226 men) with brain MRI and anthropometric measures (i.e. the full sample). A subsample (n=2,703) had body MRI measures available. We excluded participants with known diagnosis of cancers, selected traumas, neurological, psychiatric, substance abuse, cardiovascular, liver, or severe infectious conditions (Note S1), with incomplete demographic or clinical data, or who withdrew their consent. We did not exclude based on common metabolic syndrome or lifestyle factors, but adjust for these in the analyses.

UK biobank has IRB approval from North West Multi-center Research Ethics Committee and its Ethics Advisory Committee (https://www.ukbiobank.ac.uk/ethics) oversees the UK biobank Ethics & Governance Framework.^17^ We obtained access to the UK biobank cohort through Application number 27412. The study was approved by the Regional Committees for Medical and Health Research Ethics (https://helseforskning.etikkom.no), and conducted in accordance with the Helsinki Declaration.

### Demographic and clinical data

We extracted demographic and clinical data (age, sex, ethnicity), and variables reflecting cardiovascular risk (including history of diabetes, hypercholesterolemia, hypertension, current cigarette smoker, current alcohol consumption), and body composition (waist/hip circumference, weight, height). We computed BMI and WHR (Table 1; Note S2).

**Table 1:** Demographics

|  | **Men**<br>**n=9226** | **Women**<br>**n=10,104** | **$\chi^2$-test / t-test**<br>**/ Wilcoxon**<br>**rank-sum test** | **p-value** |
| --- | --- | --- | --- | --- |
| Age (year)<sup>1</sup> | 62.5±7.5 | 62.1±7.2 | 4.5 | **7.1×10<sup>-6</sup>** |
| European ancestry, N (%)<sup>2</sup> | 8891 (96.4) | 9822 (97.2) | 10.7 | **0.0011** |
| Smoker, N (%)<sup>2,3</sup> | 397 (4.3) | 298 (2.9) | 25.1 | **5.4×10<sup>-7</sup>** |
| Alcohol drinker, N (%)<sup>2,3</sup> | 8751 (94.9) | 9444 (93.5) | 16.5 | **5.0×10<sup>-5</sup>** |
| Height (cm)<sup>1</sup> | 176.3±6.6 | 162.7±6.2 | 147.2 | **0** |
| Weight (kg)<sup>1</sup> | 83.2±13.1 | 68.3±12.6 | 80.4 | **0** |
| BMI<sup>1</sup> | 26.7±3.8 | 25.8±4.6 | 15.5 | **4.5×10<sup>-54</sup>** |
| Waist circumference (cm)<sup>1</sup> | 93.4±10.3 | 82.2±11.4 | 71.6 | **0** |
| Hip circumference (cm)<sup>1</sup> | 100.7±7.1 | 100.7±9.6 | 0.1 | 0.8895 |
| WHR<sup>1</sup> | 0.9±0.1 | 0.8±0.1 | 120.6 | **0** |
| Diabetes, N (%)<sup>2,3</sup> | 169 (1.8) | 84 (0.8) | 36.6 | **1.5×10<sup>-9</sup>** |
| Hypercholesterolemia, N (%)<sup>2,3</sup> | 1178 (12.8) | 734 (7.3) | 163.3 | **2.2×10<sup>-37</sup>** |
| Hypertension, N (%)<sup>2,3</sup> | 1998 (21.7) | 1531 (15.2) | 136.3 | **1.8×10<sup>-31</sup>** |

*Notes:* Report mean ± standard deviation for continuous variables, unless otherwise stated. P-values<0.05 considered significant. *Abbreviations:* BMI – body mass index; WHR – waist-to-hip ratio.
<sup>1</sup> Welch two sample t-test.
<sup>2</sup> $\chi^2$ -test.
<sup>3</sup> Self-reported.

### MRI acquisition

Participants underwent 3T brain and 1·5T body/liver MRI on the same day and site. Brain MRI was available from three sites (Cheadle, Reading, and Newcastle), while body/liver MRI was from one site (Cheadle). Similar scanners/protocols were used across all sites^17^ (Note S3).

### MRI processing

We processed brain MRI DICOM images *in-house* using FreeSurfer (version 5·3·0; http://www.freesurfer.net). We extracted mean cortical thickness and white surface area from the cortical parcellation, and volumes of cortical/cerebellum gray/white matter, brain stem, CSF, lateral ventricle, third ventricle, thalamus, hippocampus, amygdala, accumbens, caudate, putamen, and pallidum. For bilateral measures, we computed the average across the hemispheres. Additionally, we extracted Euler numbers as a proxy of image quality.

We extracted body and liver MRI data processed for abdominal and liver fat, and thigh muscle volumes. Body MRI was processed for visceral adipose tissue (VAT), abdominal subcutaneous adipose tissue (ASAT), total abdominal adipose tissue (VAT+ASAT), and total thigh muscle volume (TTMV) by AMRA (https://www.amramedical.com). Liver MRI was processed for liver proton density fat fraction (PDFF) by Perspectum Diagnostics (https://perspectum-diagnostics.com/) (Note S4).

### MRI quality control

Among the 33,303 participants with available brain MRI, 21,395 met the inclusion criteria. Of these, we excluded two participants with missing Euler numbers prior to iteratively excluding Euler outliers defined as participants with higher negative Euler numbers that exceeded three standard deviations from the mean in either hemisphere. We iterated until no outliers remained, resulting in eight iterations. This led to further exclusion of 2063 participants, yielding a total sample of n=19,330 (Figure S1; Note S5).

For body MRI, participants labelled with severe motion artifacts, corrupted data, broken coil element, outer field-of-view inhomogeneities, or metal contamination were removed. We further excluded participants with incomplete measures. In total, 857 participants were excluded, largely due to missing liver PDFF measures (n=676), yielding a total body MRI subsample of n=2,703 (Figure S1).

### Statistical analysis

We investigated the sample demographics across and within sexes. Categorical variables were compared using χ^2^-test. For normally/non-normally distributed continuous variables we used the two-sample t-test/Wilcoxon rank-sum test. For unequal variance across sexes, t-test was replaced by Welch approximation. Normality was assessed by visual evaluation of density plots (Figure S2-S3). For brain structure, we assessed density plots for expected distribution patterns (Figure S4-S5), and scatter plots of body-brain associations (Figures S6-S13).

For descriptive purposes, and to establish fundamental properties of the dataset, we initially assessed age- and sex-related associations on body composition and brain structure using a three-step multiple linear regression approach: *model 1a* included age, age^2^, and sex; *model 1b* additionally included age-by-sex and age^2^-by-sex interactions; and *model 1c* additionally included metabolic/lifestyle variables, including ethnicity (relates to ethnic differences in adipose tissue distribution/accumulation^10^), current cigarette smoking (yes/no), current alcohol consumption (yes/no), diabetes (yes/no), hypertension (yes/no), and hypercholesterolemia (yes/no). Model 1c was only applied to body composition measures.

In the main analyses, building upon model 1b/c, we tested for linear and quadratic associations between brain structure (dependent variable) and body composition, specifically (i) anthropometric measures, and (ii) regional/specific body MRI measures. We used a three-step multiple linear regression approach: *model 2a* included linear body composition term; *model 2b* additionally included a quadratic body composition term; and *model 2c* additionally included metabolic/lifestyle variables. Model 2a/b extends model 1b, while model 2c extends model 1c.

We used regression models of incremental complexity to motivate the fully adjusted models, and to explore the importance of nonlinearities in body-brain association. Separate analyses were conducted for the full sample and body MRI subsample (when applicable). For brain MRI, we additionally adjusted for intracranial volume (except cortical thickness), image quality (average Euler number), and site (when applicable). Categorical variables were included as factors, continuous variables were mean-centered.

We evaluated model residuals for normality using residual vs fitted value and Q-Q plots, leading to log-transformation of dependent variables for models showing a significant departure from normality, namely: all outcomes for sample description analyses of body composition, and CSF, lateral, and third ventricle volumes for brain MRI analyses (Figures S14-S19 presents selected illustrations). Remaining dependent variables were not log-transformed.

All statistical analyses were conducted in R (version 3·5·2; https://www.r-project.org). We used *lm* for the regression analyses (Note S6 presents pseudocode), and computed the partial correlation coefficients, *r*, effect size directly from the t-statistics for continuous variables and via Cohen’s d for categorical variables.^18^ We used Bonferroni correction to adjust for multiple comparison at α=0·05 across N independent tests, defined as N=(3+8)(1+17)+17=215, which is the number of regression models from sample description and main analyses for both the full sample and body MRI subsample, counting partly overlapping models once. This resulted in a study-wide significance threshold of *p*≤α/N=0·0002. We present the overall global picture of significant findings from *model 2c*, and the range of *p*-values and *r* effect sizes (absolute values; indicated by |*r*|). The full results are presented in the supplemental material.

## Results

### Demographic variables

The full subject sample (n=19,330) included more women (n=10,104; 52·3%) than men (n=9226; 47·7%), and was largely of self-identified European ancestry (96·8%). The age-range was 44-82 years. Compared to women, men had significantly higher age, more alcohol consumers and cigarette smokers, higher anthropometric measures (except hip circumference), and more were diagnosed with diabetes, hypercholesterolemia, and hypertension (Table 1; Figure S2). Similarly, the body MRI subsample (n=2,703) included more women (n=1,496; 55·4%) than men (n=1,207; 44·7%), the age-range was 46-77 years, and men had higher age and more adverse factors than women (Table S1; Figure S3).

Primarily for descriptive purposes, the sample characteristics were further explored for age- and sex-effects on body composition and brain structure in Note S7.

### Brain structure and anthropometric measures

Analyses in the full sample revealed a robust body-brain association pattern, and we observed overall negative and nonlinear associations between anthropometric body composition measures and global brain volumes, and positive associations with accumbens (Figures 1, S6-S8). Using model 2c (Figure 2a), the largest negative effect sizes were observed for cerebellum gray matter, brain stem, and cortical gray matter volumes, mean surface area, and cortical thickness (Figure 2b), together with significant quadratic terms suggestive of negative and accelerating reductions for cerebellum gray matter, brain stem, and cortical gray matter volume and thickness (Figure 2c) with higher anthropometric measures. We also observed decreasing cortical and cerebellum white matter volumes with increasing body composition measures, with indications of accelerating differences for cerebellum white matter volume. There were positive associations between anthropometric measures and CSF, third ventricle, and lateral ventricle. Lateral ventricle also displayed positive quadratic associations suggestive of accelerated expansion. Subcortical structures exhibited mixed results. Higher anthropometric measures were negatively associated with pallidum, and hippocampus, and positively associated with amygdala, accumbens, and putamen. There were significant negative quadratic terms for thalamus, hippocampus, amygdala, and accumbens, suggestive of accelerating reductions for thalamus and hippocampus, while amygdala and accumbens increases were quadratically attenuated. The effect sizes of the quadratic terms were small (|*r*|≤0·06). Findings did not reach significance across all anthropometric measures.

**Figure 1:**
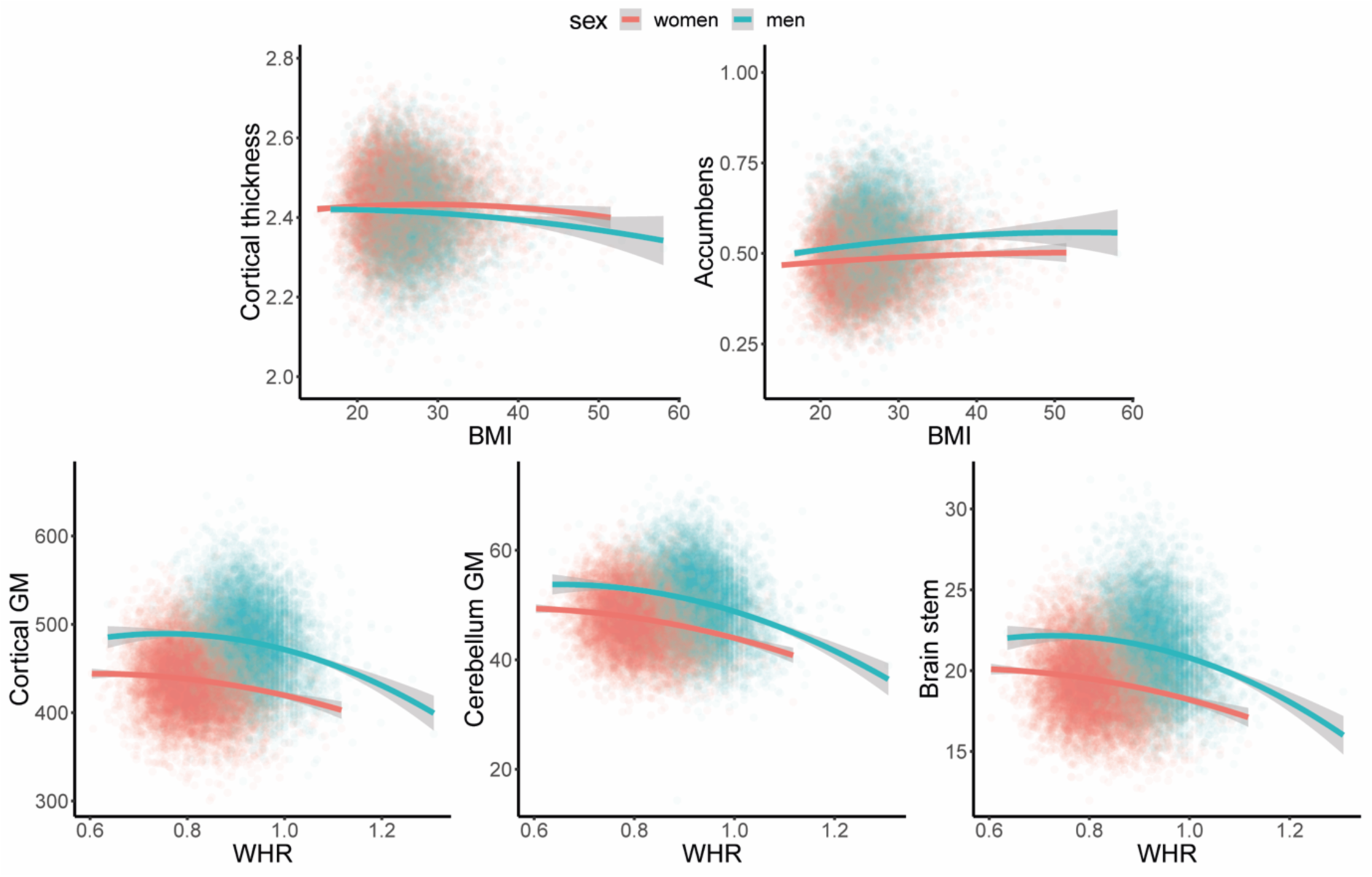
Association pattern between selected brain structures and body composition measures. *Notes:* Regression lines are modeled as brain structure = body composition + body composition^2^. The confidence intervals are indicated in gray. Illustrations are split on sex (commonly a significant factor in neuroimaging studies), but are not adjusted for other confounders. *Abbreviations:* BMI – body mass index; WHR – waist-to-hip ratio.

**Figure 2:**
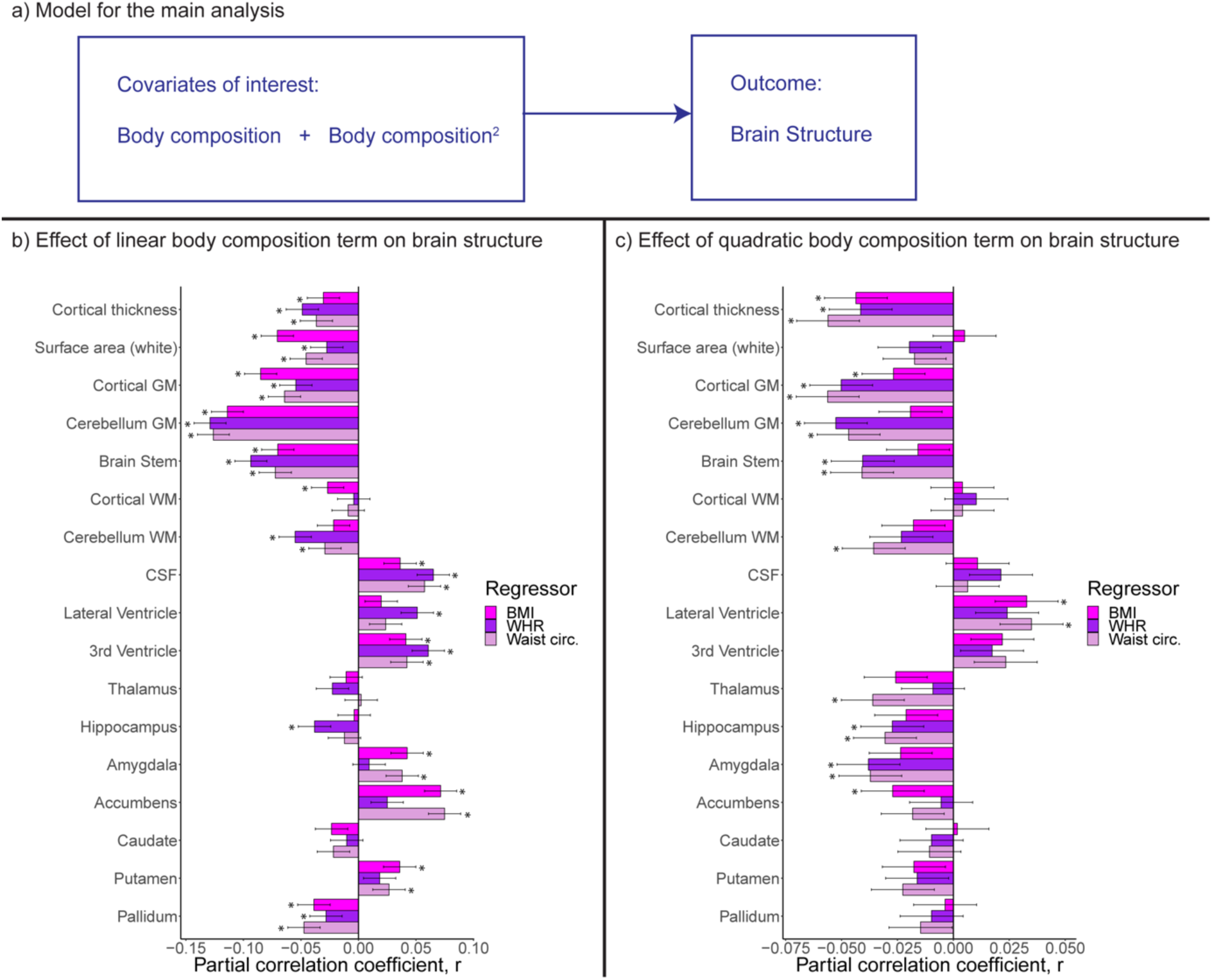
Body-brain association patterns in healthy individuals (n=19,330). *Notes:* Results from model 2c (Panel a) that investigates body-brain connections through the inclusion of linear (Panel b) and quadratic (Panel c) body composition terms after adjusting for lifestyle/metabolic factors. Additional confounding variables included age, age^2^, sex, age-by-sex, age^2^-by-sex, intracranial volume (except cortical thickness), Euler number, and site. Significant associations indicated by *. Dependent variables CSF, lateral/3^rd^ ventricle were log-transformed. *Abbreviations:* BMI – body mass index; circ. – circumference; GM – gray matter; WHR – waist-to-hip ratio; WM – white matter.

These findings were robust across models 2a/b/c, with some adjustment of significant levels and effect sizes, with *p* in [2·8×10^−80^, 0·0002], and |*r*| in [0·03, 0·14] (Figure S20; Tables S5-S7). For model 2c, compared to models 2a/b, we observed attenuation of significance levels and effect sizes with *p* in [1.6×10^−72^, 0·0002] and |*r*| in [0·03, 0·13] between body and brain anthropometrics. Additionally, we here observed significant associations (|*r|* in [0·03, 0·07]) between self-reported diabetes, hypertension, or hypercholesterolemia diagnosis, and brain structure, including lower cortical and cerebellum white and gray matter volumes, and higher ventricular volumes. Current cigarette smoking was associated with lower mean cortical thickness, while current alcohol consumption was not significantly associated with brain structure.

### Brain structure, anthropometric, and body MRI measures

Analyses in the body MRI subsample (Figure 3a) revealed significant negative associations between BMI and surface area and cortical gray matter volume, and between WHR and caudate volume. For body MRI metrics (Figure 3b), liver PDFF was negatively associated with cortical gray matter thickness/volume, while TTMV was positively associated with accumbens. The associations were similar across model 2a/b/c, with anthropometric measures showing significant *p* in [3.0×10^−06^, 0·0002], and |*r*| in [0·07, 0·09], and regional body MRI measures showing significant *p* in [1·3×10^−07^, 8·9×10^−05^], and |*r*| in [0·08, 0·1]. There were fewer significant findings for anthropometric measures in model 2c compared to 2a/b (Figures S21-S22; Tables S8-S15).

**Figure 3:**
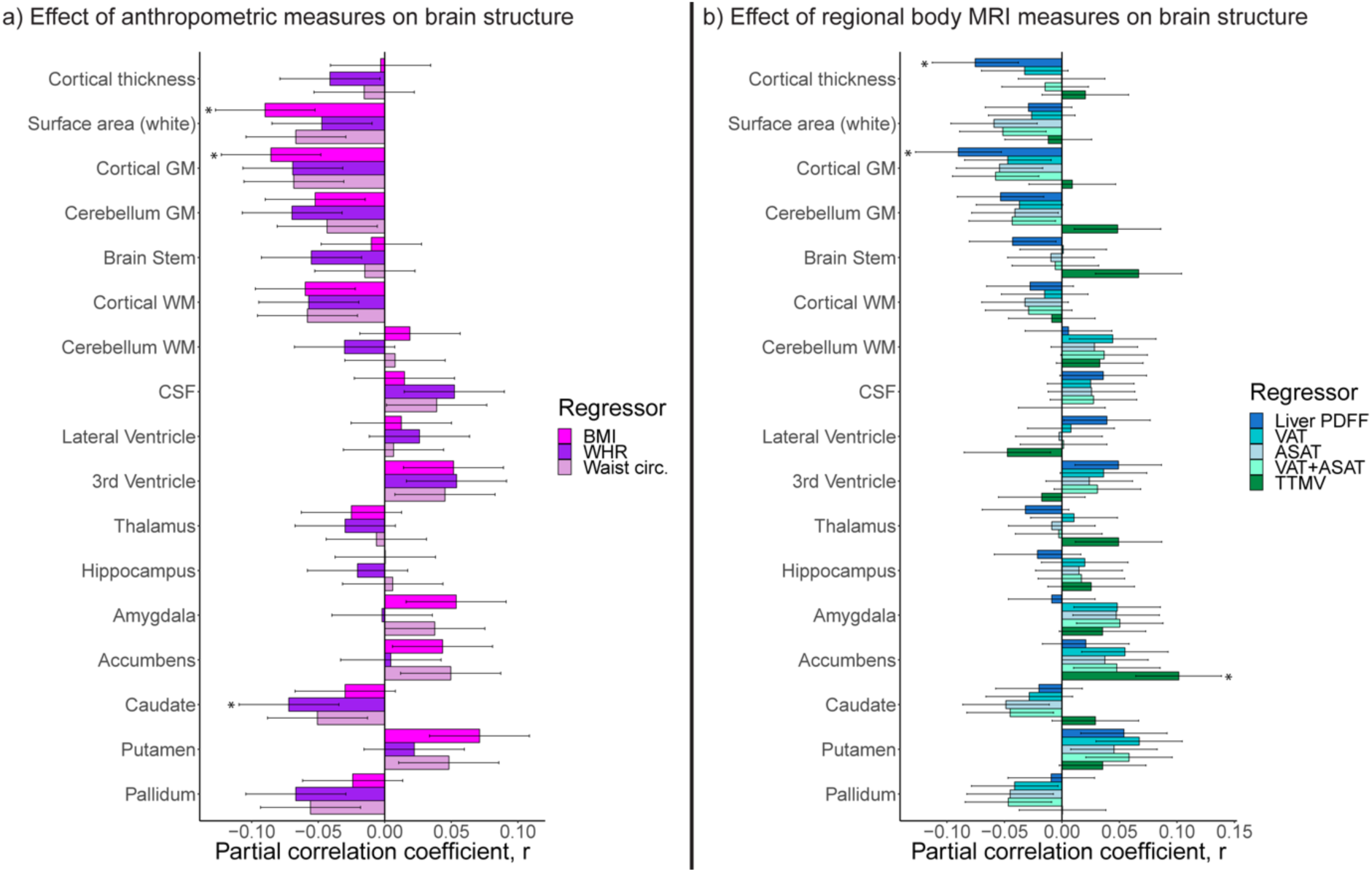
Linear body-brain association pattern in healthy individuals for the MRI subsample (n=2,703). *Notes:* Results from model 2c that investigates body-brain connections through the inclusion of linear and quadratic body composition terms after adjusting for lifestyle/metabolic factors, for a) anthropometric body composition measures, and b) regional body MRI measures on brain structure, on brain structure. Significant associations indicated by *. Dependent variables CSF, lateral/3^rd^ ventricle were log-transformed. *Abbreviations:* ASAT – abdominal subcutaneous adipose tissue; BMI – body mass index; circ. – circumference; GM-gray matter; PDFF – proton density fat fraction; TTMV – total thigh muscle volume; VAT – visceral adipose tissue; WHR – waist-to-hip ratio; WM – white matter.

## Discussion

In the largest study of body-brain connections to date, we mapped normal body-brain connections in 19,330 healthy participants without secondary disease effects. We identified a comprehensive picture of small, but highly significant, effects across body composition and brain structures, linking normally varying anthropometric measures to the majority of included brain structures. Further, we examined the novel body MRI measures (n=2,703) in this context. We revealed significant associations between liver fat and cortical gray matter volume and thickness, and between thigh muscle volume and accumbens volume.

For global brain measures, we observed the strongest, negative and nonlinear, associations between anthropometric body composition measures and cerebellar and cortical gray matter, and brain stem structures. These findings suggest accelerating cortical/cerebellum and brain stem reductions with increasing body composition levels, but also of attenuation at low body composition. Furthermore, we observed positive associations for CSF, lateral, and third ventricles. These findings are in line with our hypothesis, and with prior research showing lower total gray matter volume,^5–8^ and regional cortical and cerebellar reductions^9^ in obese individuals. Brain atrophy is observed at higher ages.^11^ Although cross-sectional evidence and small effects, our findings suggests accelerating brain atrophy also at higher adiposity levels, possibly relating to regulatory differences in brain and body lipids. Prior studies on subcortical structures and obesity are limited. Here, we showed a mixed subcortical association pattern, that were similar across anthropometric measures. Accumbens, a structure associated with motivation and reward and part of the dopamine motivation system,^19^ showed the strongest positive associations. The association between accumbens volume and body composition is generally in line with a previous study documenting larger accumbens volume in children with increased genetic risk for obesity,^20^ and supports the assumption of a critical role of brain mechanisms for reward and reinforcement learning for lifestyle and dietary choices and obesity.

Higher liver fat was associated with lower/thinner cortical gray matter volume/thickness in healthy individuals. Non-alcoholic fatty liver disease is linked with metabolic factors, show increasing prevalence rate similar to that of obesity, and increased mortality from cardiovascular disease, and patients benefit from weight loss lifestyle interventions.^21^ Our findings point towards ongoing biological processes related to liver fat that may affect brain structures, or vice versa, in healthy individuals. We may only speculate that lifestyle interventions for addressing non-alcoholic fatty liver disease may be important also for brain health.

Contrary to our expectations, we did not observe any associations between visceral adipose tissue and brain structures. Instead, we observed a largely analogous pattern across anthropometric and regional body composition measures. Effect sizes were generally stronger for anthropometric measures, which may reflect combinatorial effects of multiple factors that likely influence these measures. The novel body MRI measures may capture more specific features, but were available for less than 15% of the sample.

Earlier studies have linked vascular risk factors to brain structure.^5,8^ Our study further corroborates this in healthy individuals, and indicates an association between fat distribution and brain structure. Effect sizes and significance levels were attenuated when we adjusted for lifestyle factors and metabolic syndrome, suggestive of complex body-brain mechanisms. Self-reported diagnosis of hypercholesterolemia, hypertension, or diabetes – factors related to metabolic health - were associated with several brain structures, while current cigarette smoking was associated with thinner cortical thickness. Thus, cardiometabolic risk factors appear important for brain health.

The observed body-brain connections cut across several body compositions measures and brain structures, and appeared fairly global. Causal mechanisms are unknown and likely highly complex and multifactorial, and our findings could be mediated by modifiable lifestyle choices with known links to obesity^1^; e.g. metabolic factors could influence both somatic and brain health, impaired brain health could influence somatic health, or the effects could be reciprocal as previously implied for obesity and depression.^2^ Physical fitness and activity – common lifestyle interventions - are associated with reduced risk for obesity,^1^ may counteract a genetic predisposition for obesity,^12^ have neuroprotective effects on the brain,^22^ and have been positively related to brain structure.^13^ We found significant associations between thigh muscle volume and accumbens, and negative nonlinear association pattern between anthropometric measures and cortical, cerebellar gray matter, and brain stem structures. Although it is premature to conclude, from a public health perspective, this nonlinear association in healthy participants may imply that lifestyle interventions for normalizing body fat composition could affect biological processes related to brain function and disease.

The high degree of somatic comorbidity in mental and other brain disorders requires a better understanding of the complex biological interactions between body and brain, and of how they relate to lifestyle or environmental factors. The observed body-brain connections could be associated with obesity-related neuroinflammatory processes.^23^ A recent large-scale meta-analysis showed increased risk for vascular dementia – similar to other vascular conditions - in both underweight and obese individuals,^24^ which is interesting in light of our observed nonlinear associations between body composition and brain structure. Shared genetic underpinnings could influence both adipose tissue accumulation and brain structure, similar to the genetic overlap relating immune-related disorders,^25^ BMI,^26^ or cardiovascular risk factors,^27^ to brain disorders. Yet other complex biological and possibly polygenic processes, lifestyle/environmental factors, and/or their combinatorial effects could influence the findings. To understand both nature and nurture of somatic comorbidities in brain disorders, and brain disorders per se, further mechanistic investigations are warranted. The findings of this study may suggest that body composition is an important confounding factor that should be considered in future case-control studies.

Small effects are common across heterogeneous psychiatric disorders and research domains, including neuroimaging and genetics, making simple underlying causal mechanisms unlikely.^28^ Combinatorial complex mechanisms of additive small effects are more likely, but these are not well captured by smaller studies prone to both false positive and negative findings.^16^ Instead, large-scale investigations are needed, where effect size convergence and increasing accuracy is obtained,^16^ but this is challenging to achieve. To our knowledge, this is the first study of its magnitude investigating body composition, fitness metrics, and brain structure in healthy individuals. Through such large-scale investigations we may better understand small, but normal, body-brain processes. This will provide us with a better understanding of putative interactive or confounding factors in psychiatric and other brain disorders, thereby enhancing our conceptual understanding of the complex mechanisms at play.

This study had some limitations. At the time of MRI, the available diagnostic information was self-reported. We did not investigate the cumulative or unit effect of alcohol consumption or cigarette smoking, nor severity of hypertension, hypercholesterolemia, or diabetes diagnosis. Liver fat was assessed from selected regions of interest and not the whole liver. The observed liver-brain associations could be influenced by alcohol consumption although this was not captured by the current study. We limited our investigations to coarser brain measures. Findings deviated somewhat from prior studies using partly overlapping samples,^7,8,29^ probably due to differences in inclusion/exclusion criteria, sample size, image processing, and analyses pipeline. The exploratory cross-sectional design makes it difficult to disentangle cause from effect, determine underlying body-brain mechanisms, and draw final conclusions from the findings. Further investigations of sex-specific body-brain trajectories, and of combinatorial or additive effects of body composition, additional metabolic markers, and fitness metrics on brain structure, together with links to cognition, sex, age, disease risk, lifestyle, and environmental factors are needed.

Strengths of the study include an unprecedented sample size, including ≥160% larger sample than prior UK biobank studies,^7,8,29^ that were assessed using standardized procedures and MRI protocols.^17^ Fully automated data cleaning, inclusion and exclusion criteria, and quality control limits chance for subjective variations or errors. We build upon confirmatory analyses of known age- and sex-related associations on both body composition^10^ and brain structure^11,30^ that largely mirrored the current knowledge, which strengthened our confidence in the reported body-brain patterns. We applied rigorous diagnosis-based exclusion criteria to capture normative body-brain associations in healthy individuals, and rigorous correction for multiple comparisons.

### Conclusions

Through large-scale body-brain mapping we link normally varying body composition measures to brain structure in a healthy population. The results imply correlated effects of adipose tissue and poor metabolic health on brain structure, affecting global brain structures, brain cavities, and accumbens, at higher measured body composition - yet the causal mechanisms remain unknown. It is of vital importance to investigate the underlying complex body-brain pathways, shared mechanisms of cardiometabolic risk factors and brain disorders, and lifestyle-related modifying factors. If brain structure alterations can be linked to lifestyle-related body composition characteristics, then preventive public health interventions for normalizing cardiometabolic risk factors, could prevent the development of disorders of the body and the brain.

## Data sharing

The UK biobank resource is open for eligible researchers upon application (http://www.ukbiobank.ac.uk/register-apply/).

## Code availability

We made use of publicly available resources for processing the image data and for conducting the statistical analyses. The project R-scripts will be made publicly available upon publication.

## Acknowledgements

The Research Council of Norway (#223273; #276082); South-Eastern Norway Regional Health Authority (#2017112); European Union’s Horizon2020 Research and Innovation Programme (CoMorMent project; Grant #847776) & European Research Council (ERC) StG (Grant #802998). This work was performed on *Services for sensitive data* (TSD), University of Oslo, Norway, with resources provided by UNINETT Sigma2 - the National Infrastructure for High Performance Computing and Data Storage in Norway.

## Authors’ contributions

*Study design:* OAA, TPG, LTW. *Data preparation and image processing:* TPG, TK, LTW. *Data management:* TPG, TK, DvdM, OBS, AW. *Analytical strategy:* DA, OF, TPG, UKH, KO, TM, IES, LTW, JL, ODL, RS. *Statistical analysis:* DA, OF, TPG, LTW. *Figures:* TPG. *Data* i*nterpretation*: OAA, TPG, KO, IES, NES, LTW. *Manuscript preparation:* OAA, OF, TPG, TK, KO, IES, NES, LTW. *Funding:* OAA, LTW. All authors revised the manuscript and approved the final version.

