## Supplemental material for "Population-based body-brain mapping links brain morphology and body composition"

**Table of contents**

**SUPPLEMENTAL FIGURES** 3

**Figure S1: Inclusion/exclusion pipeline, including overview of automatic quality control step.** 3

**Figure S2: Distribution continuous demographic variables for the whole sample (n=19,330).** 4

**Figure S3: Distribution of continuous demographic variables for the body MRI subsample (n=2703).** 5

**Figure S4: Distribution of included brain structures for the whole sample (n=19,330).** 6

**Figure S5: Distribution of included brain structures for the body MRI subsample (n=2703).** 7

**Figure S6: Association pattern between measures of brain structure and BMI (n=19,330).** 8

**Figure S7: Association pattern between measures of brain structure and WHR (n=19,330).** 9

**Figure S8: Association pattern between measures of brain structure and waist circumference (n=19,330).** 10

**Figure S9: Association pattern between measures of brain structure and liver PDFF (n=2,703).** 11

**Figure S10: Association pattern between measures of brain structure and VAT (n=2,703).** 12

**Figure S11: Association pattern between measures of brain structure and ASAT (n=2,703).** 13

**Figure S12: Association pattern between measures of brain structure and VAT+ASAT (n=2,703).** 14

**Figure S13: Association pattern between measures of brain structure and TTMV (n=2,703).** 15

**Figure S14: Residual versus fitted value plots and Q-Q plots for all body composition models with (left) and without (right) log-transformation of dependent variables (full sample; n=19,330).** 16

**Figure S15: Residual versus fitted value plots and Q-Q plots for all body composition models with (left) and without (right) log-transformation of dependent variables (body MRI subsample; n=2703).** 17

**Figure S16: Residual versus fitted value plots and Q-Q plots for all body composition models after log-transformation of CSF, lateral ventricle, and 3rd ventricle (full sample; n=19,330).** 18

**Figure S17: Residual versus fitted value plots and Q-Q plots for CSF, lateral ventricle, and 3rd ventricle with (left) and without (right) log-transformation (full sample; n=19,330).** 19

**Figure S18: Residual versus fitted value plots and Q-Q plots for all body composition models after log-transformation of CSF, lateral ventricle, and 3rd ventricle (body MRI subsample; n=2703).** 20

**Figure S19: Evaluation of multiple linear regression model residuals for normality using residual versus fitted value plots and Q-Q plots for CSF, lateral ventricle, and 3rd ventricle with (left) and without (right) log-transformation (body MRI subsample; n=2703).** 21

**Figure S20-a: Linear body-brain associations in healthy across models 2a/b/c (n=19,330).** **22**

**Figure S20-b: Quadratic body-brain associations in healthy across models 2b/c (n=19,330).** **23**

**Figure S21-a: Linear body-brain associations in healthy across models 2a/b/c (n=2703) for anthropometric measures.** **24**

**Figure S21-b: Quadratic body-brain associations in healthy across models 2b/c (n=2703) for anthropometric measures.** **25**

**Figure S22-a: Linear body-brain associations in healthy across models 2a/b/c (n=2703) for body MRI measures.** **26**

**Figure S22-b: Quadratic body-brain associations in healthy across models 2b/c (n=2703) for body MRI measures.** **27**

**SUPPLEMENTAL TABLES** **28**

**Table S1: Demographics for the body MRI subsample (n=2703).** **28**

**Tables S2-S15 are presented sheet-wise in a separate supplemental excel document.** **29**

**SUPPLEMENTAL NOTES** **30**

**Note S1: Exclusion criteria for the study** **30**

**Note S2: Extracted/computed demographic and clinical variables** **31**

**Note S3: MRI acquisition** **32**

**Note S4: Body MRI processing details** **32**

**Note S5: Brain MRI Quality control** **33**

**Note S6: Linear regression models using the *lm* in r** **34**

**Note S7: Sample description analyses of body composition and brain structure.** **35**

**REFERENCES** **38**

**Supplemental Figures**

**Figure S1: Inclusion/exclusion pipeline, including overview of automatic quality control step.**

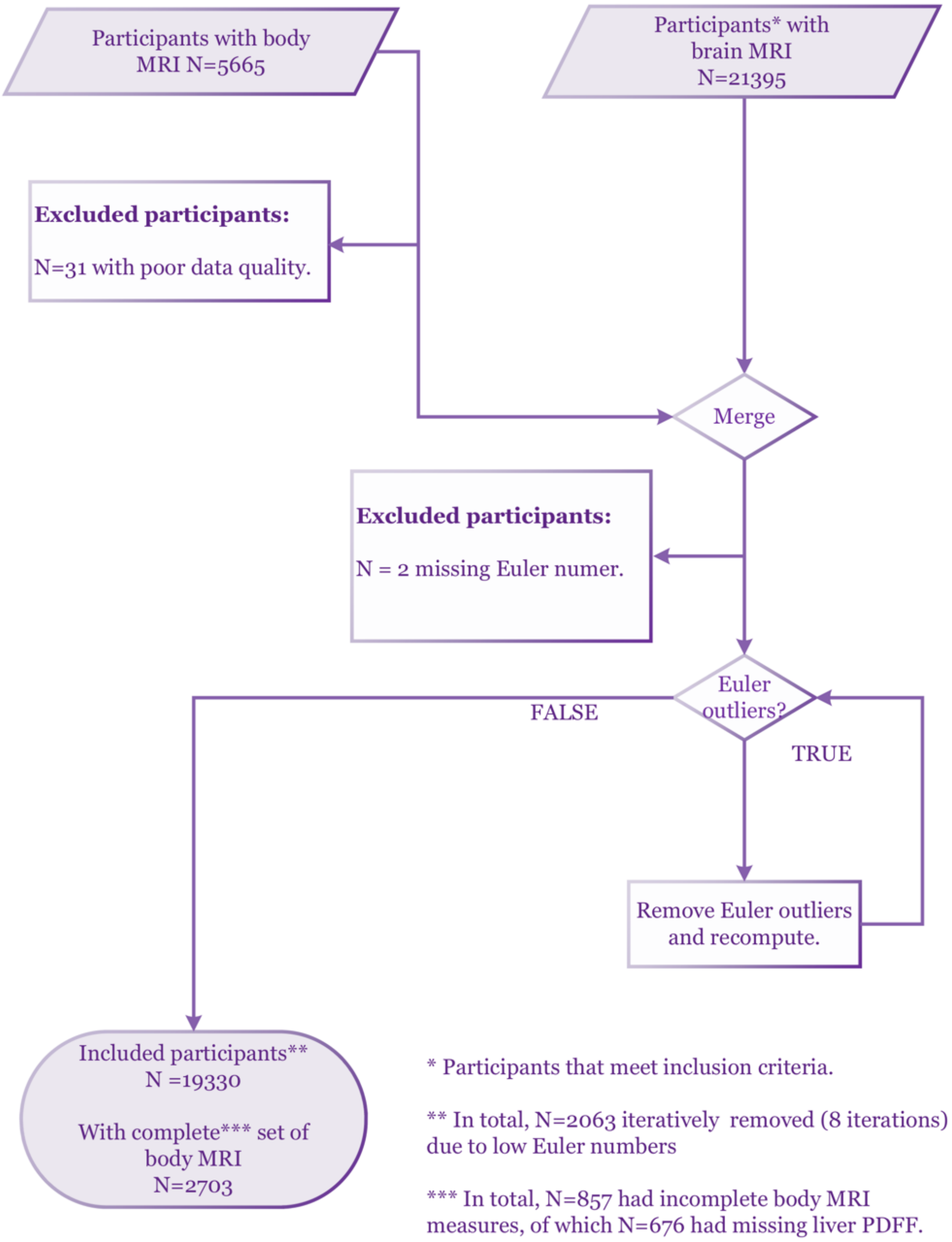


*Note:* We had brain MRI on 33,303 participants. Initially, 11,608 were excluded based on diagnosis exclusion criteria, or incomplete demographic data. *Abbreviations:* MRI – magnetic resonance imaging.

**Figure S2: Distribution continuous demographic variables for the whole sample (n=19,330).**

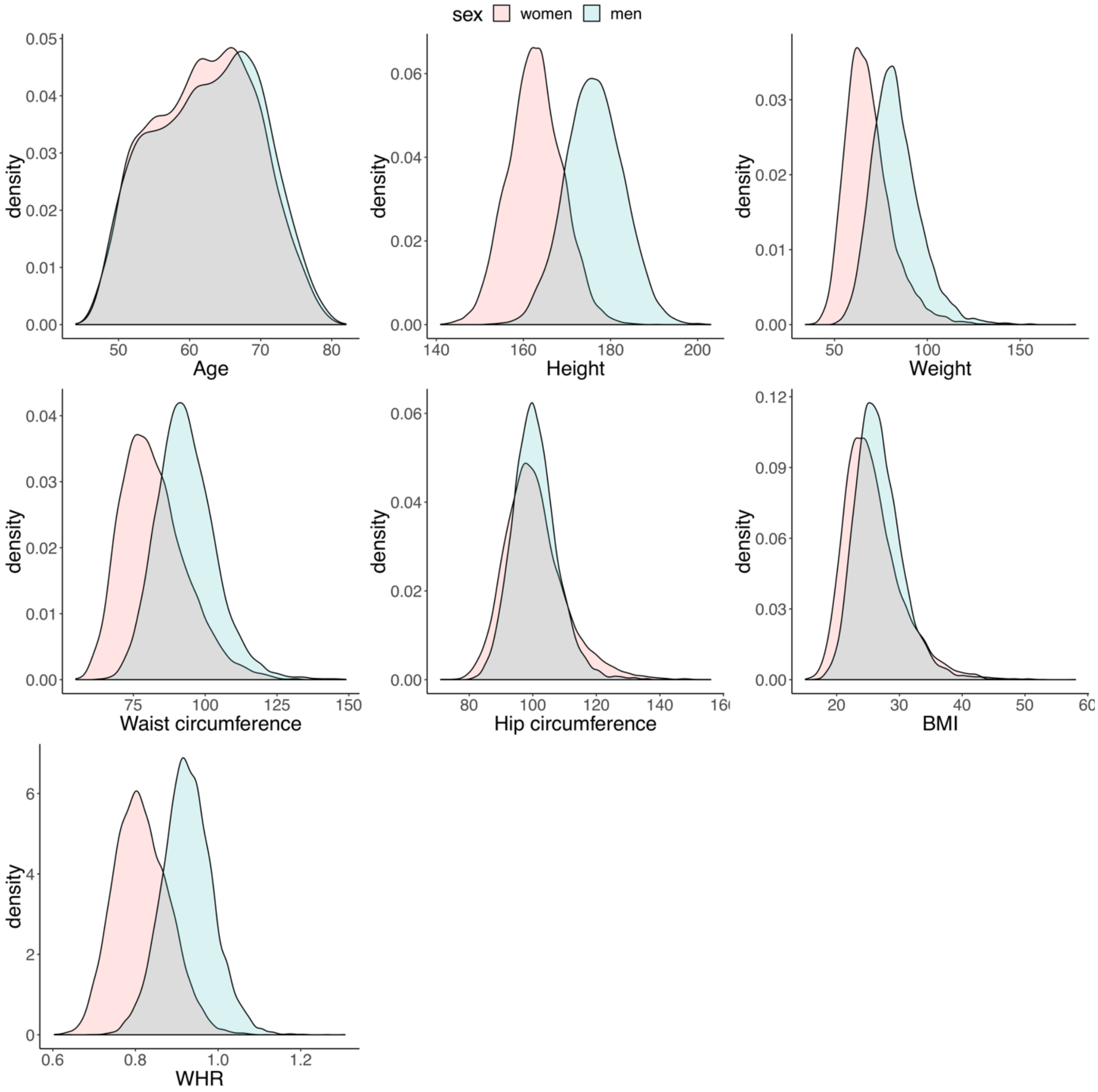


*Notes:* Weight in kg, height in cm. *Abbreviations:* ASAT – abdominal subcutaneous adipose tissue; BMI - body mass index; WHR – waist-to-hip ratio.

**Figure S3: Distribution of continuous demographic variables for the body MRI subsample (n=2703).**

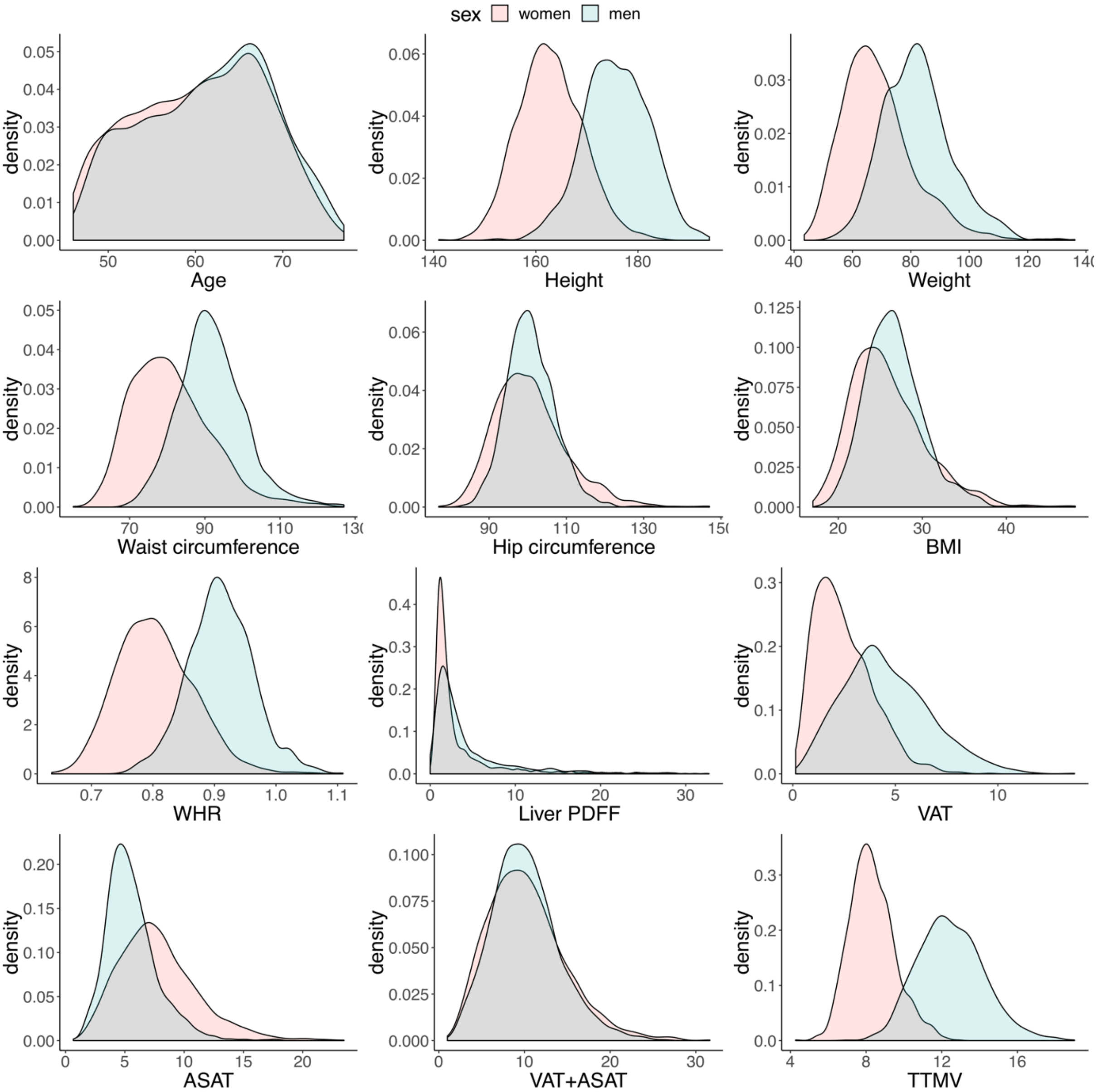


*Notes:* Weight in kg, height in cm. *Abbreviations:* ASAT – abdominal subcutaneous adipose tissue; BMI - body mass index; PDFF – Proton density fat fraction; TTMV - Total thigh muscle volume; VAT – visceral adipose tissue; VAT+ASAT – total abdominal adipose tissue; WHR – waist-to-hip ratio.

**Figure S4: Distribution of included brain structures for the whole sample (n=19,330).**

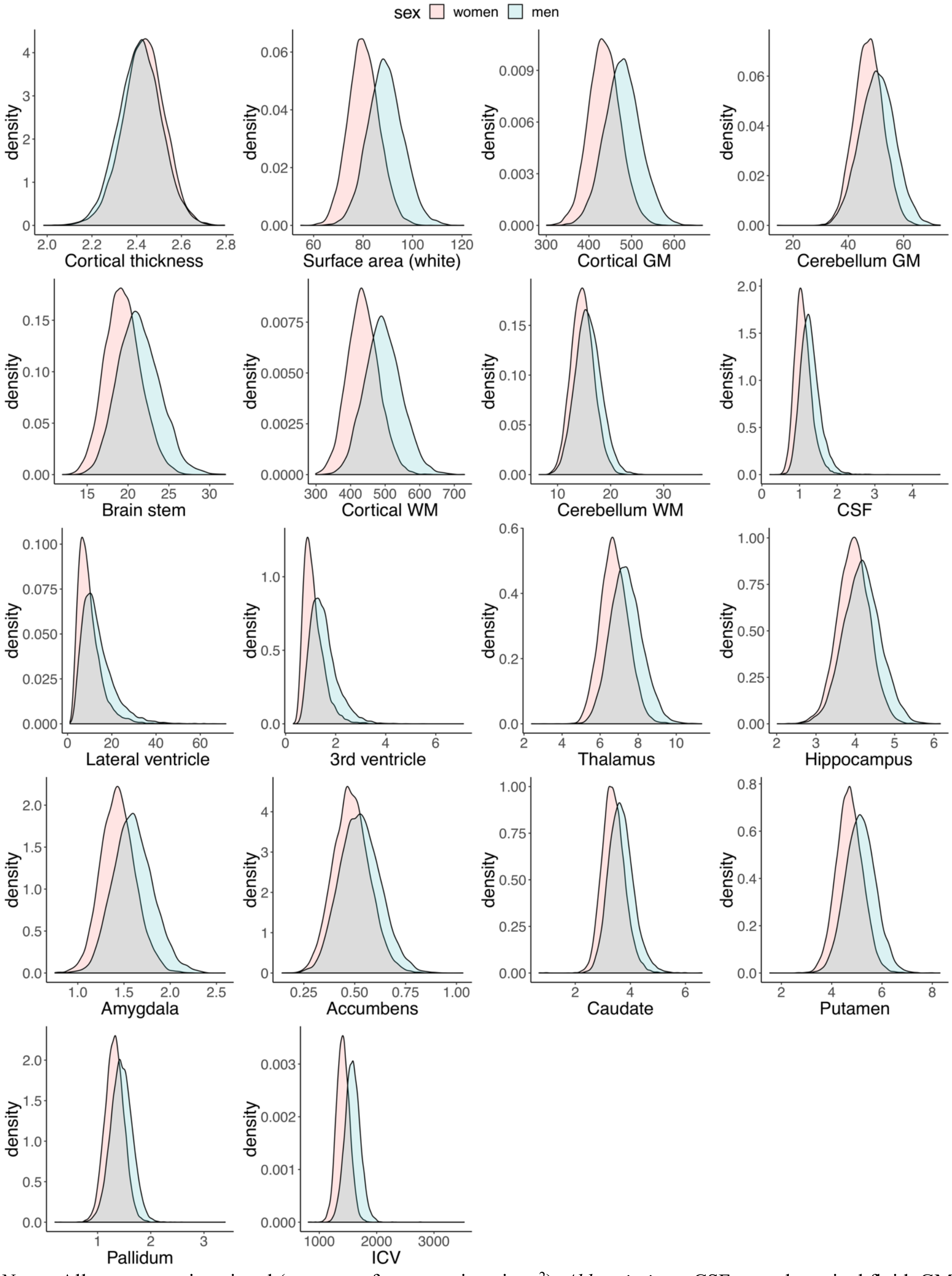


*Notes:* All structures given in ml (except surface area given in $m^2$). *Abbreviations:* CSF – cerebrospinal fluid; GM – gray matter; ICV – intracranial volume; WM – white matter.

**Figure S5: Distribution of included brain structures for the body MRI subsample (n=2703).**

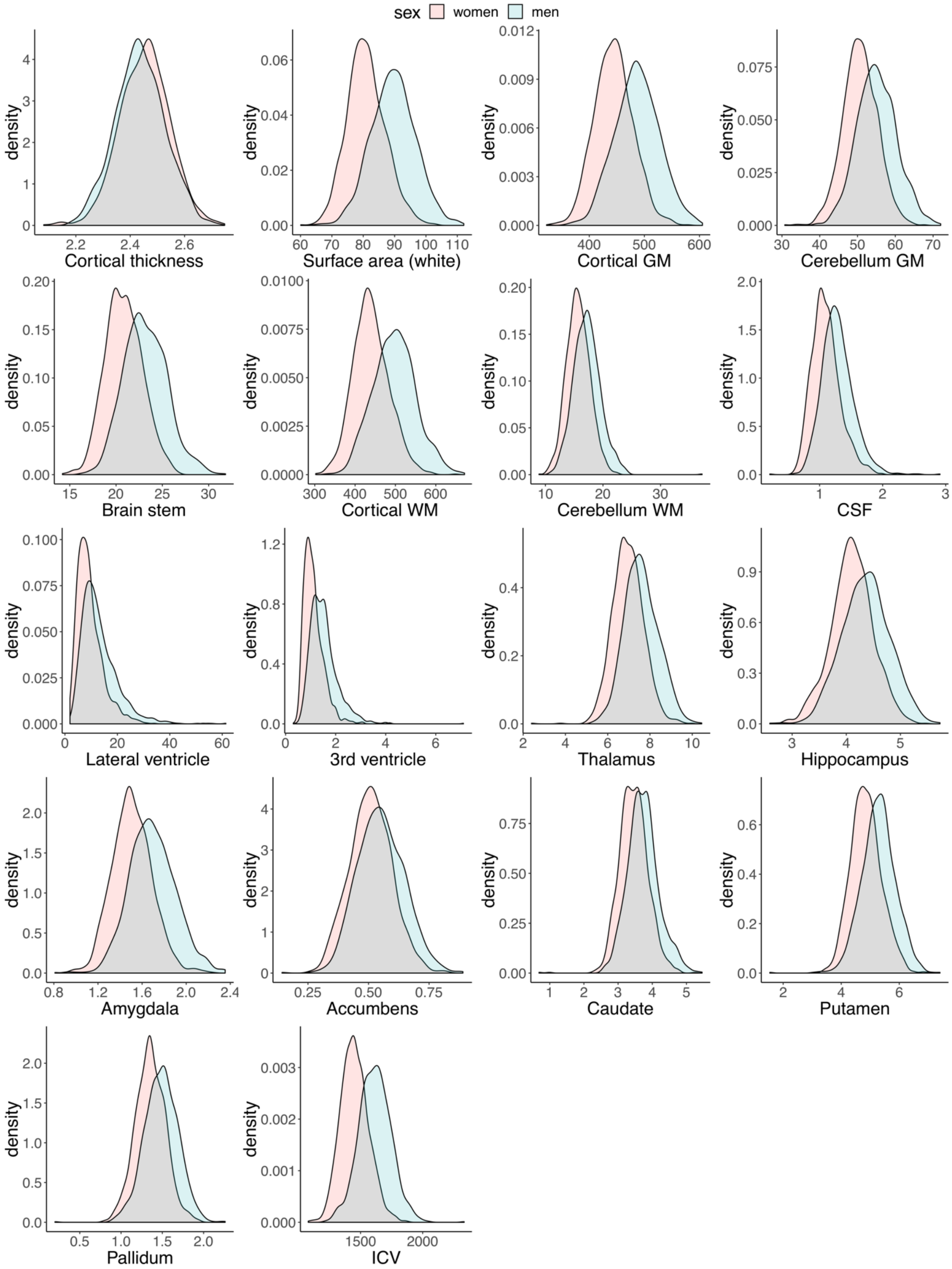


*Notes:* All structures given in ml (except surface area given in $m^2$ ). *Abbreviations:* CSF – cerebrospinal fluid; GM – gray matter; ICV – intracranial volume; WM – white matter.

**Figure S6: Association pattern between measures of brain structure and BMI (n=19,330).**

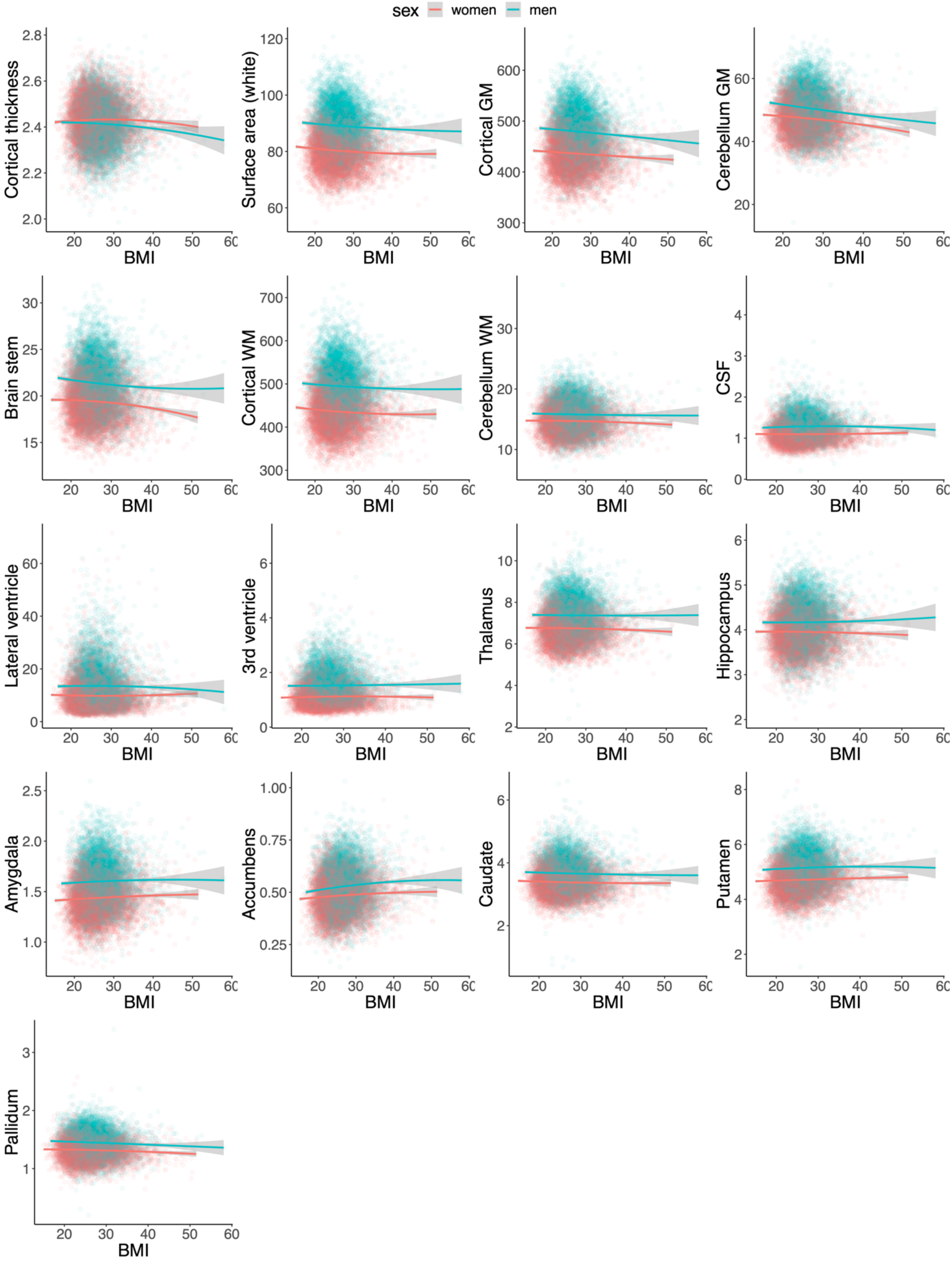


*Notes:* Regression lines are modeled as brain structure = body composition + body composition$^2$. The confidence intervals are indicated in gray. Illustrations were split on sex (commonly a significant factor in neuroimaging studies), but were not adjusted for other confounders. *Abbreviations:* BMI – body mass index.

**Figure S7: Association pattern between measures of brain structure and WHR (n=19,330).**

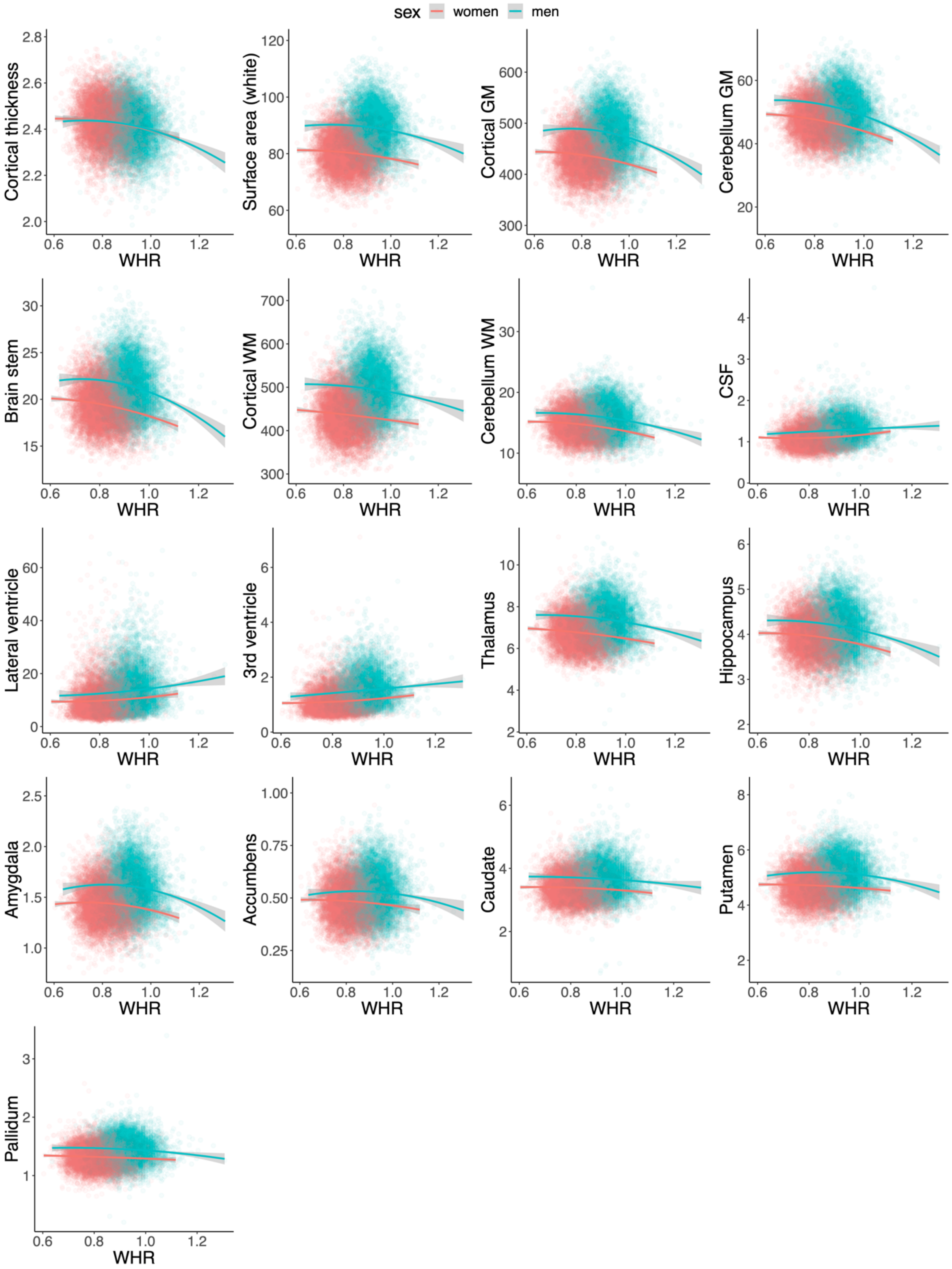


*Notes:* Regression lines are modeled as brain structure = body composition + body composition$^2$. The confidence intervals are indicated in gray. Illustrations were split on sex (commonly a significant factor in neuroimaging studies), but were not adjusted for other confounders. *Abbreviations:* WHR – waist-to-hip ratio.

**Figure S8: Association pattern between measures of brain structure and waist circumference (n=19,330).**

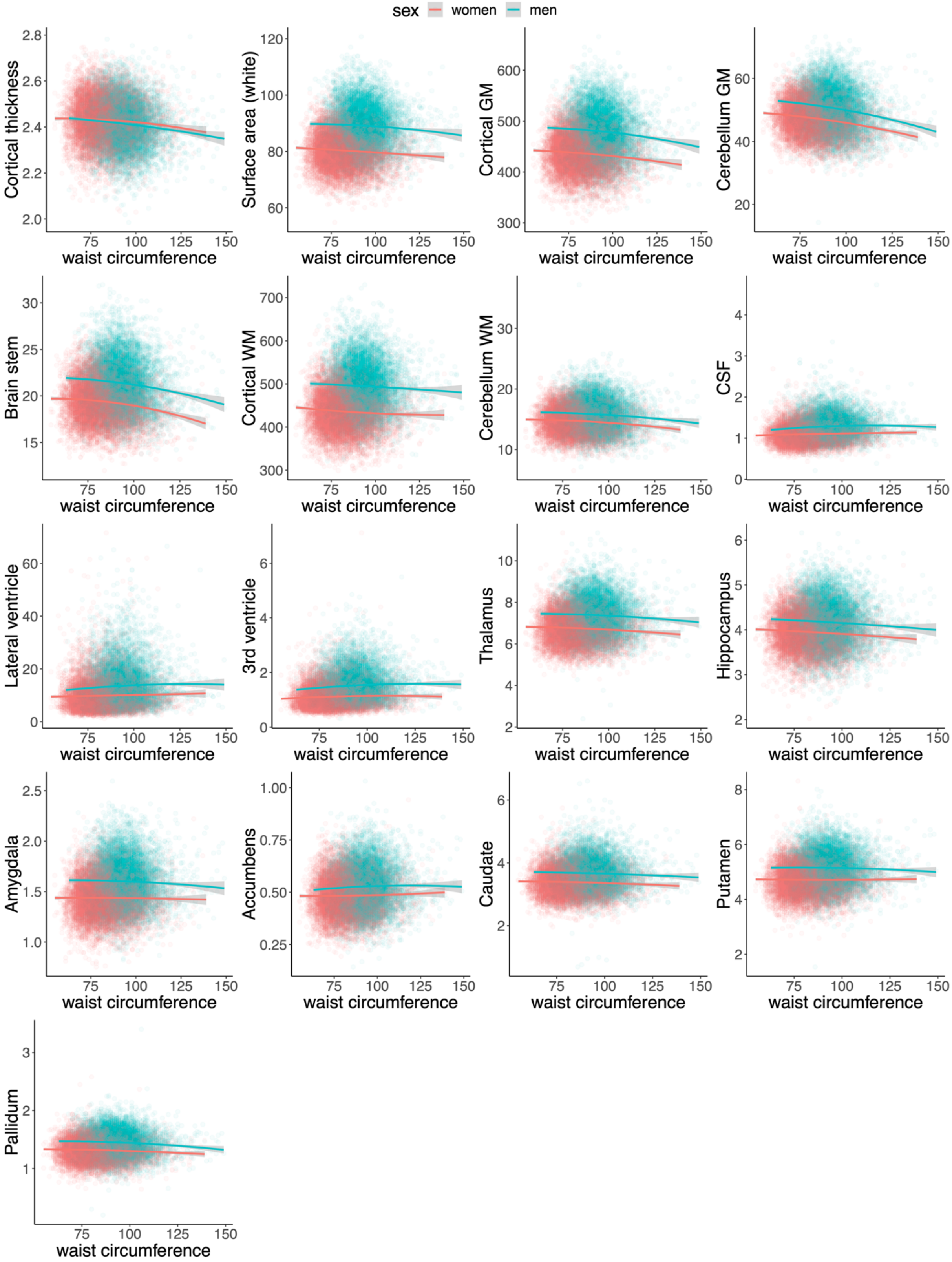


*Notes:* Regression lines are modeled as brain structure = body composition + body composition$^2$. The confidence intervals are indicated in gray. Illustrations were split on sex (commonly a significant factor in neuroimaging studies), but were not adjusted for other confounders.

**Figure S9: Association pattern between measures of brain structure and liver PDFF (n=2,703).**

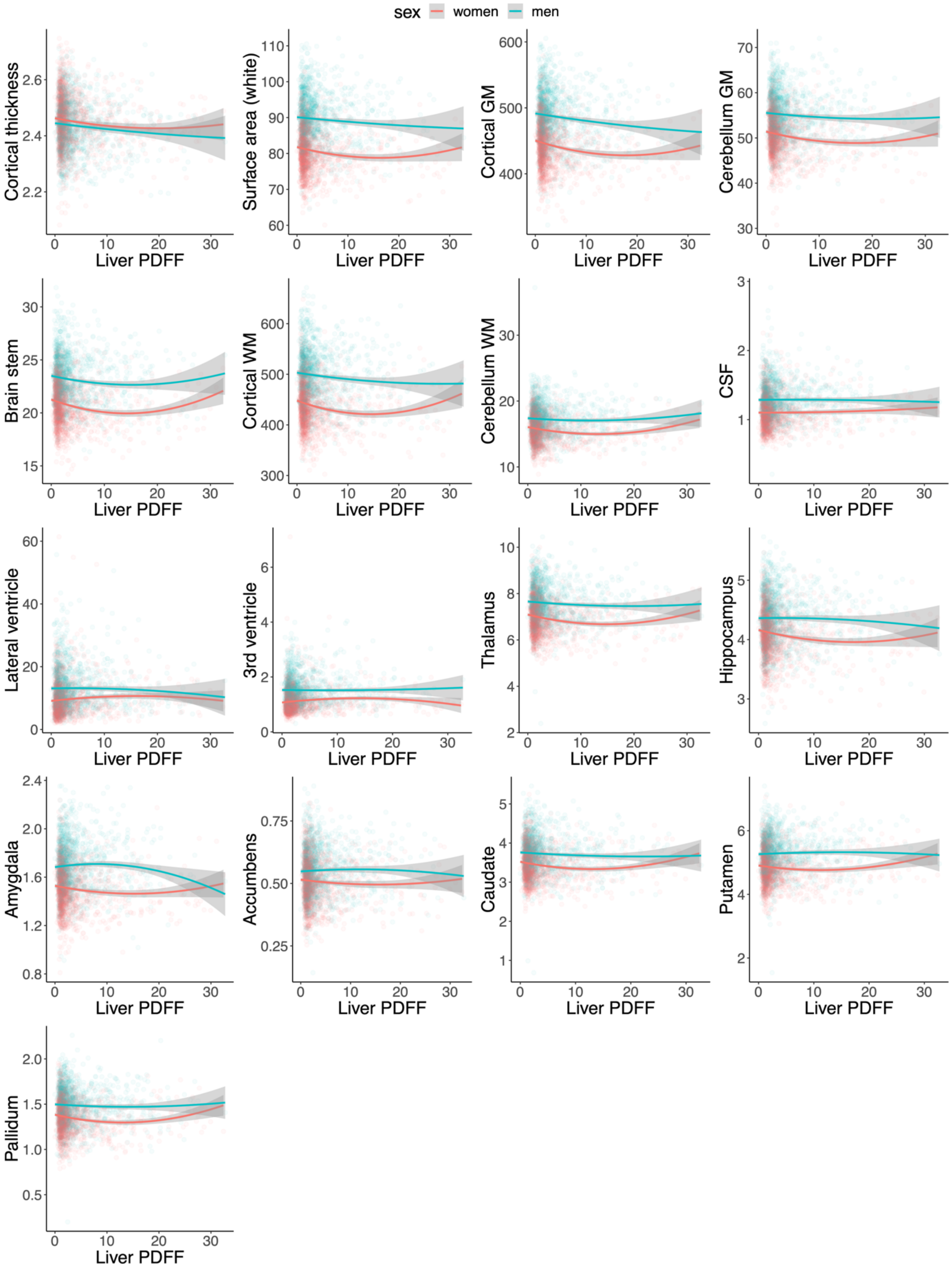


*Notes:* Regression lines are modeled as brain structure = body composition + body composition$^2$. The confidence intervals are indicated in gray. Illustrations were split on sex (commonly a significant factor in neuroimaging studies), but were not adjusted for other confounders. *Abbreviations*: PDFF – proton density fat fraction.

**Figure S10: Association pattern between measures of brain structure and VAT (n=2,703).**

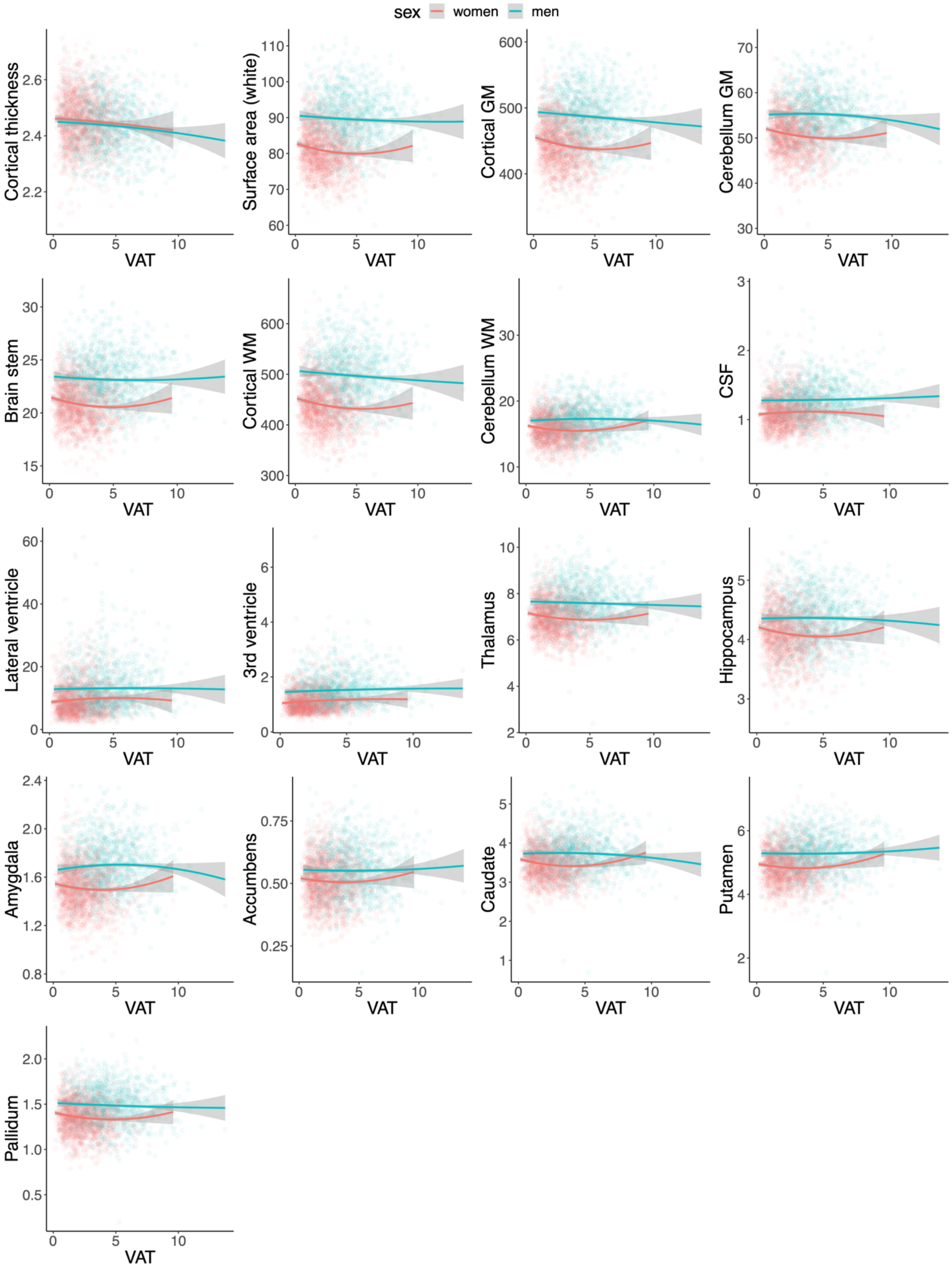


*Notes:* Regression lines are modeled as brain structure = body composition + body composition$^2$. The confidence intervals are indicated in gray. Illustrations were split on sex (commonly a significant factor in neuroimaging studies), but were not adjusted for other confounders. *Abbreviations*: VAT – visceral adipose tissue.

**Figure S11: Association pattern between measures of brain structure and ASAT (n=2,703).**

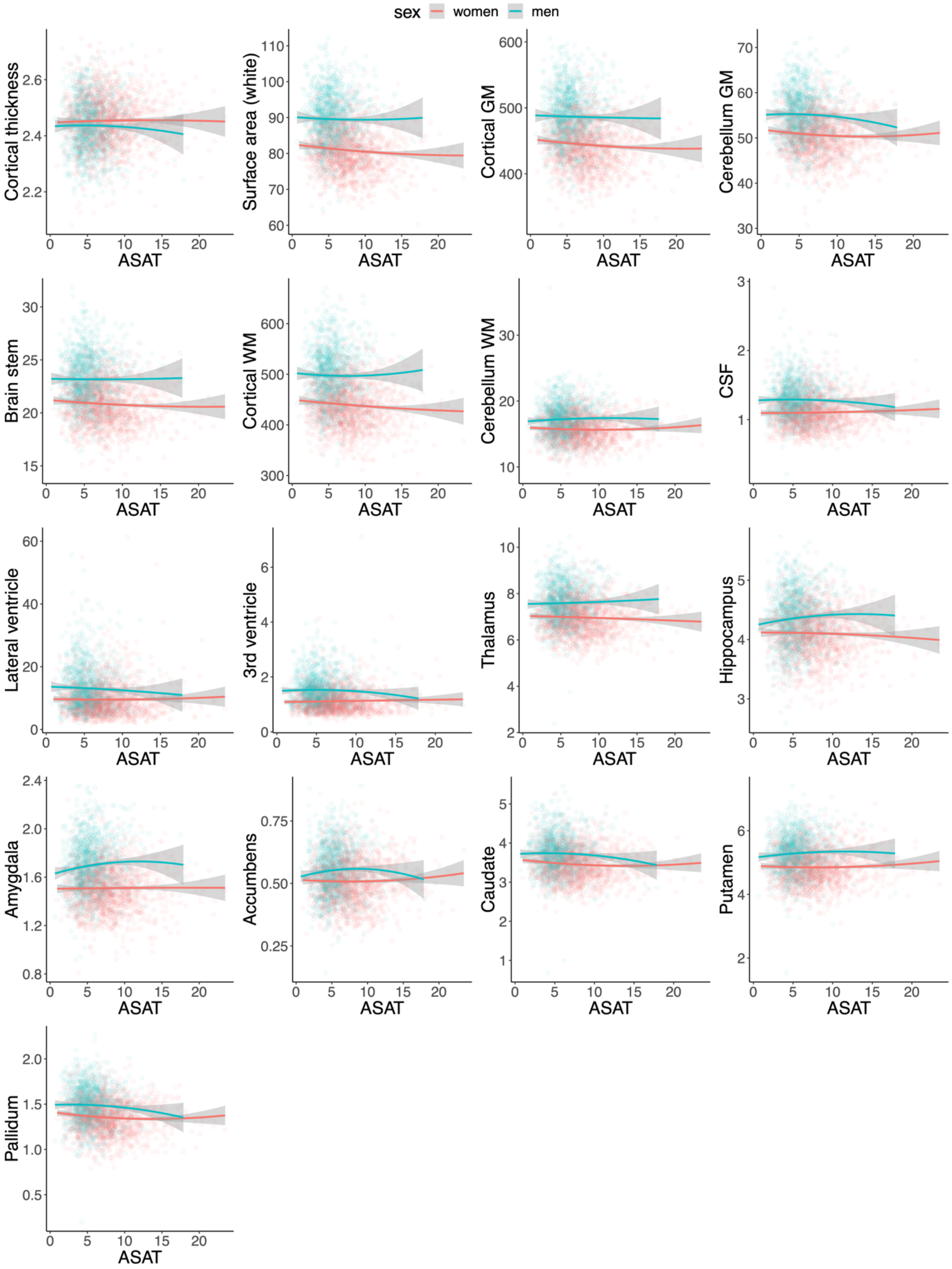


*Notes:* Regression lines are modeled as brain structure = body composition + body composition$^2$. The confidence intervals are indicated in gray. Illustrations were split on sex (commonly a significant factor in neuroimaging studies), but were not adjusted for other confounders. *Abbreviations*: ASAT – abdominal subcutaneous adipose tissue.

**Figure S12: Association pattern between measures of brain structure and VAT+ASAT (n=2,703).**

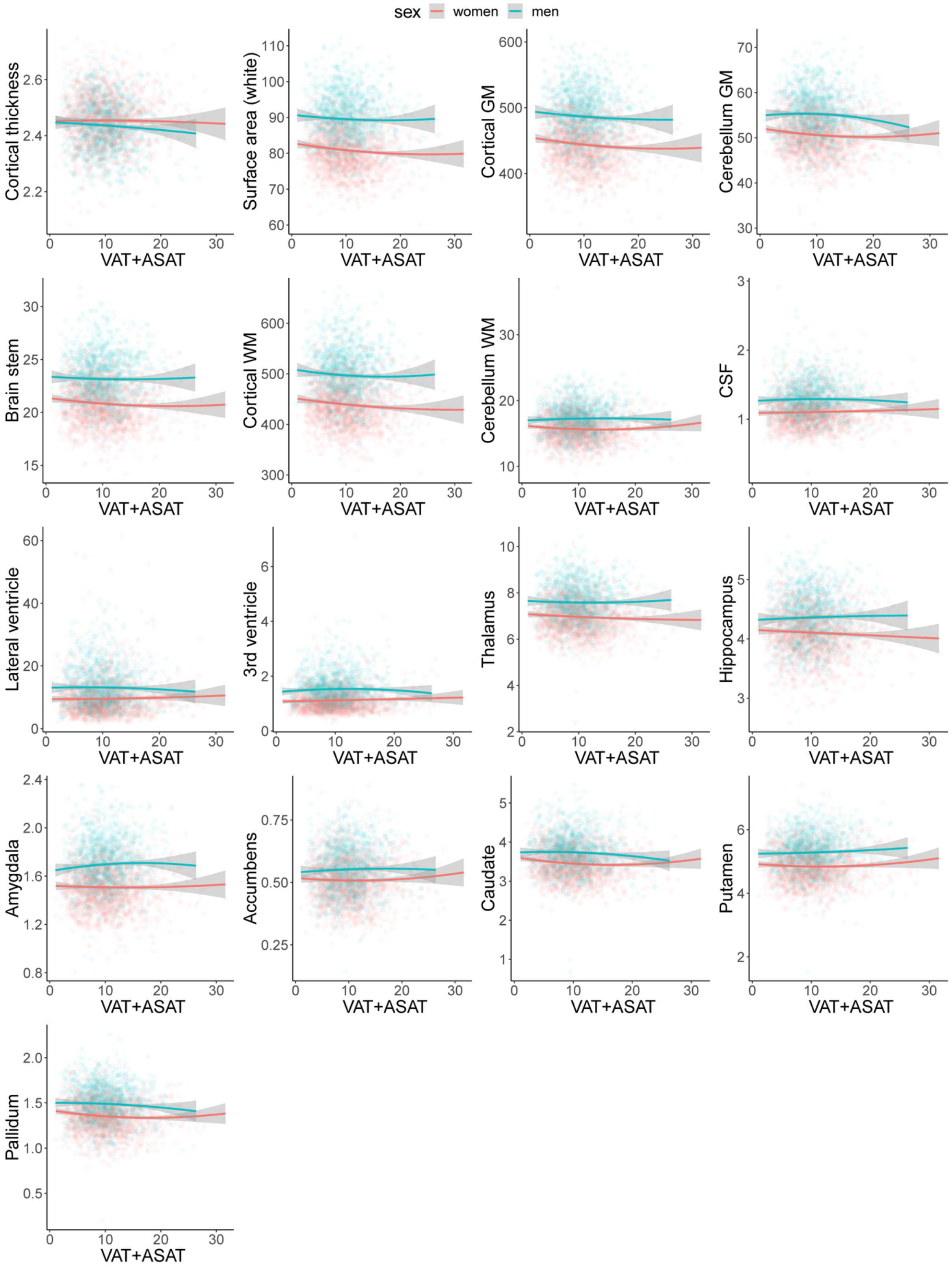


*Notes:* Regression lines are modeled as brain structure = body composition + body composition$^2$. The confidence intervals are indicated in gray. Illustrations were split on sex (commonly a significant factor in neuroimaging studies), but were not adjusted for other confounders. *Abbreviations*: VAT+ASAT – total abdominal adipose tissue.

**Figure S13: Association pattern between measures of brain structure and TTMV (n=2,703).**

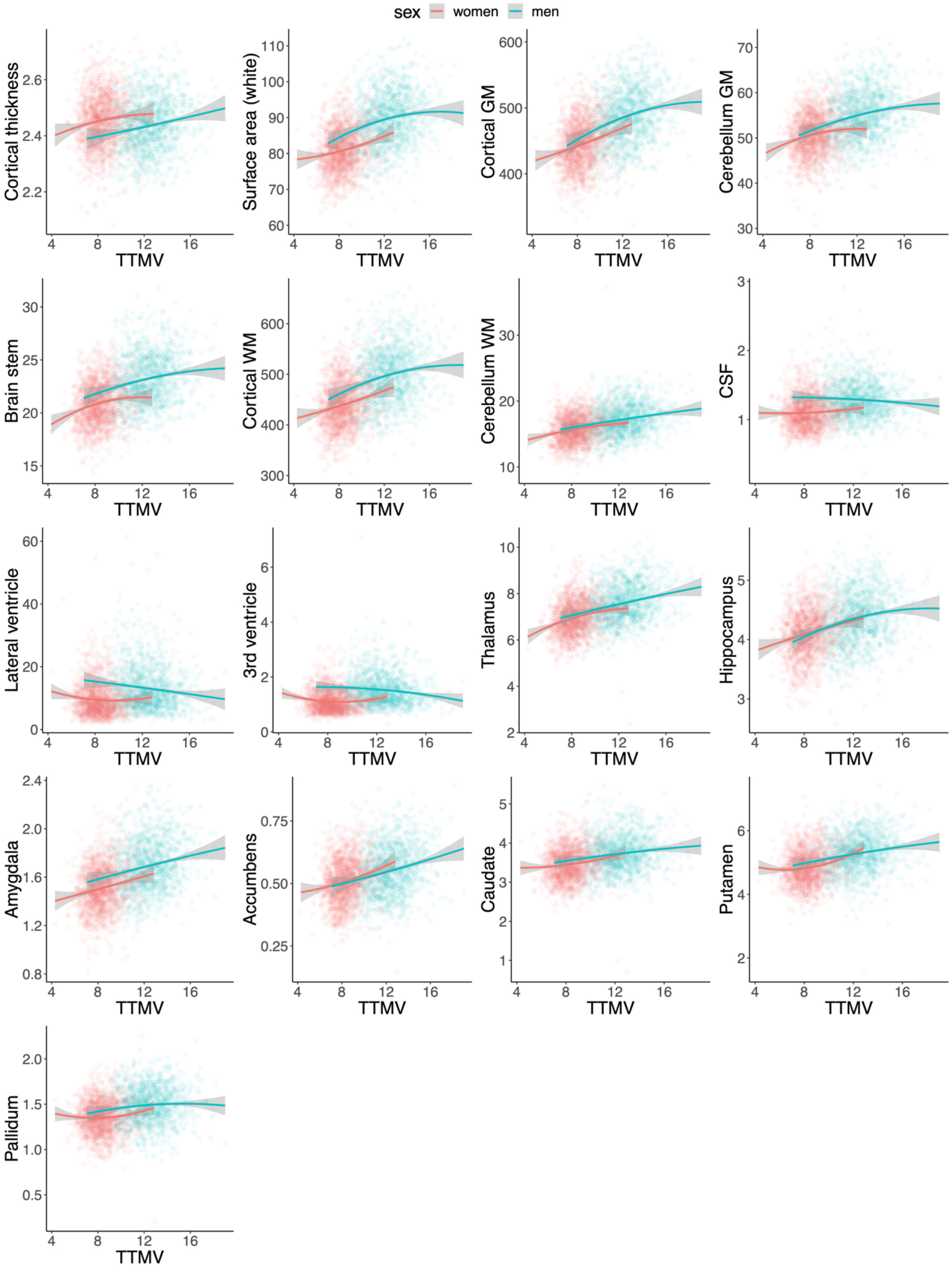


*Notes:* Regression lines are modeled as brain structure = body composition + body composition$^2$. The confidence intervals are indicated in gray. Illustrations were split on sex (commonly a significant factor in neuroimaging studies), but were not adjusted for other confounders. *Abbreviations*: TTMV – total thigh muscle volume.

**Figure S14: Residual versus fitted value plots and Q-Q plots for all body composition models with (left) and without (right) log-transformation of dependent variables (full sample; n=19,330).**

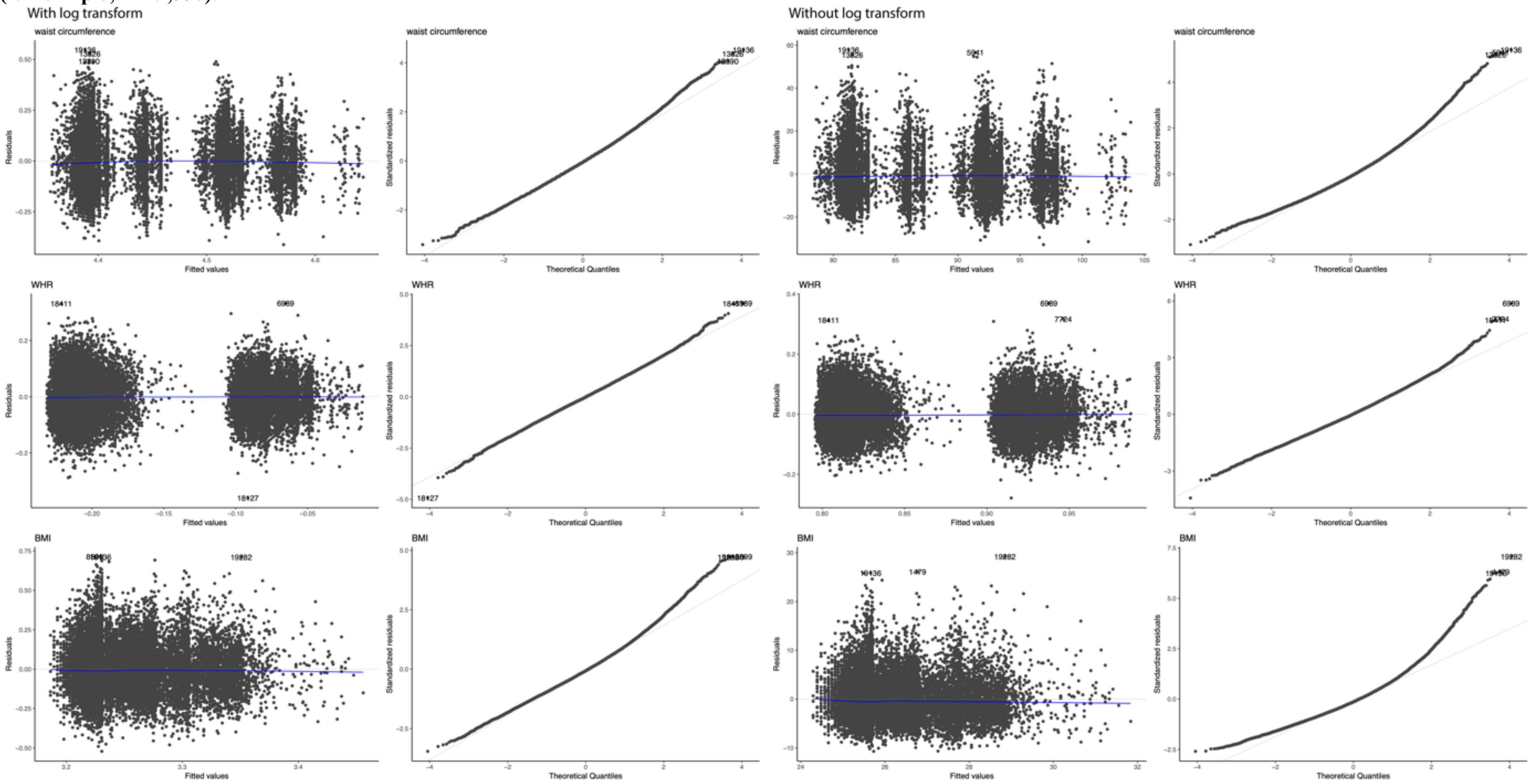


*Notes:* Residuals derived from model 1c. *Abbreviations:* BMI – body mass index; WHR – waist-hip-ratio.

**Figure S15: Residual versus fitted value plots and Q-Q plots for all body composition models with (left) and without (right) log-transformation of dependent variables (body MRI subsample; n=2703).**

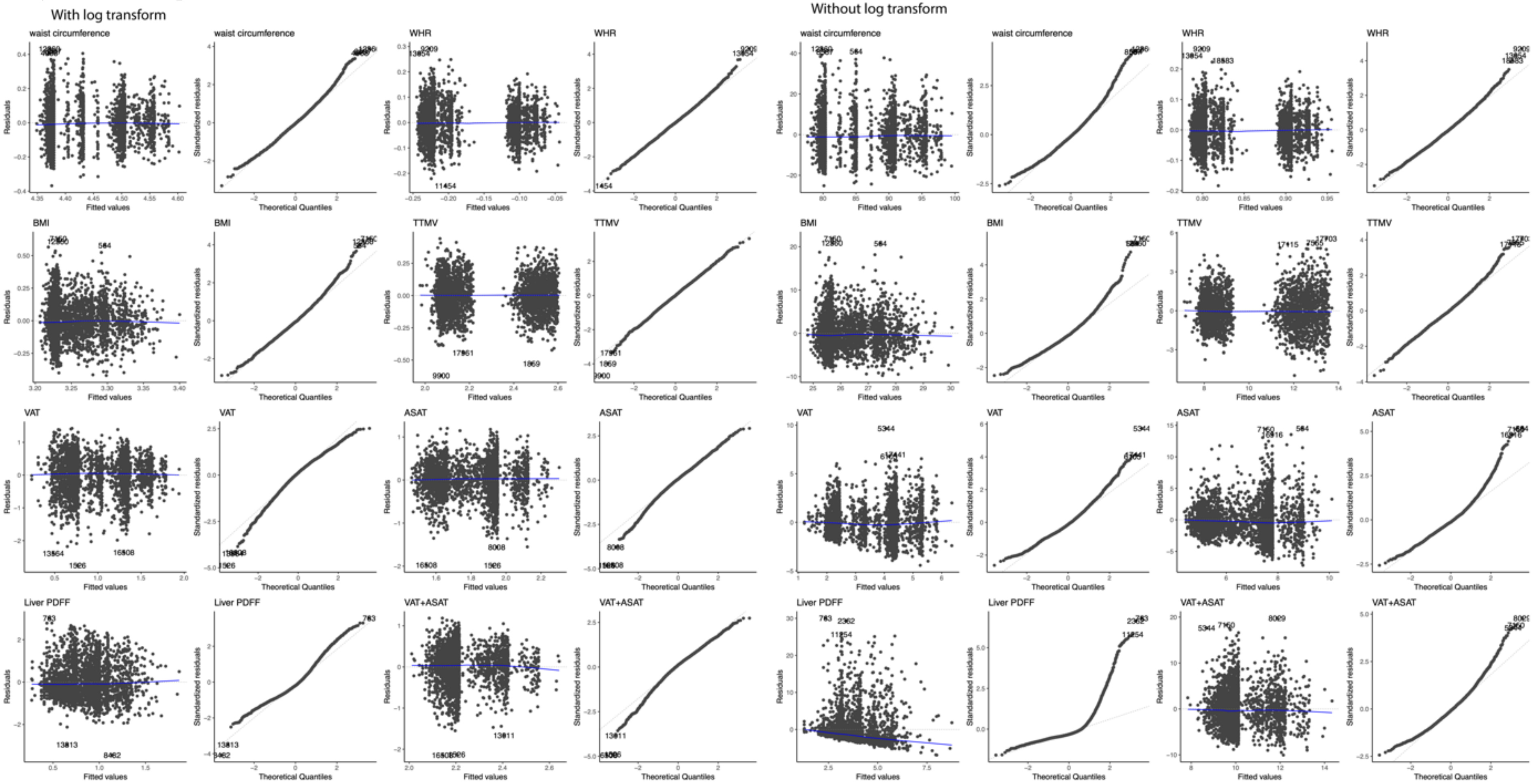


*Notes:* Residuals derived from model 1c. For two participants, we approximated zero liver PDFF measures by 0.1, smallest measure >0 was 0.23 prior to log-transformation.
*Abbreviations:* BMI – body mass index; WHR – waist-hip-ratio.

**Figure S16: Residual versus fitted value plots and Q-Q plots for all body composition models after log-transformation of CSF, lateral ventricle, and 3rd ventricle (full sample; n=19,330).**

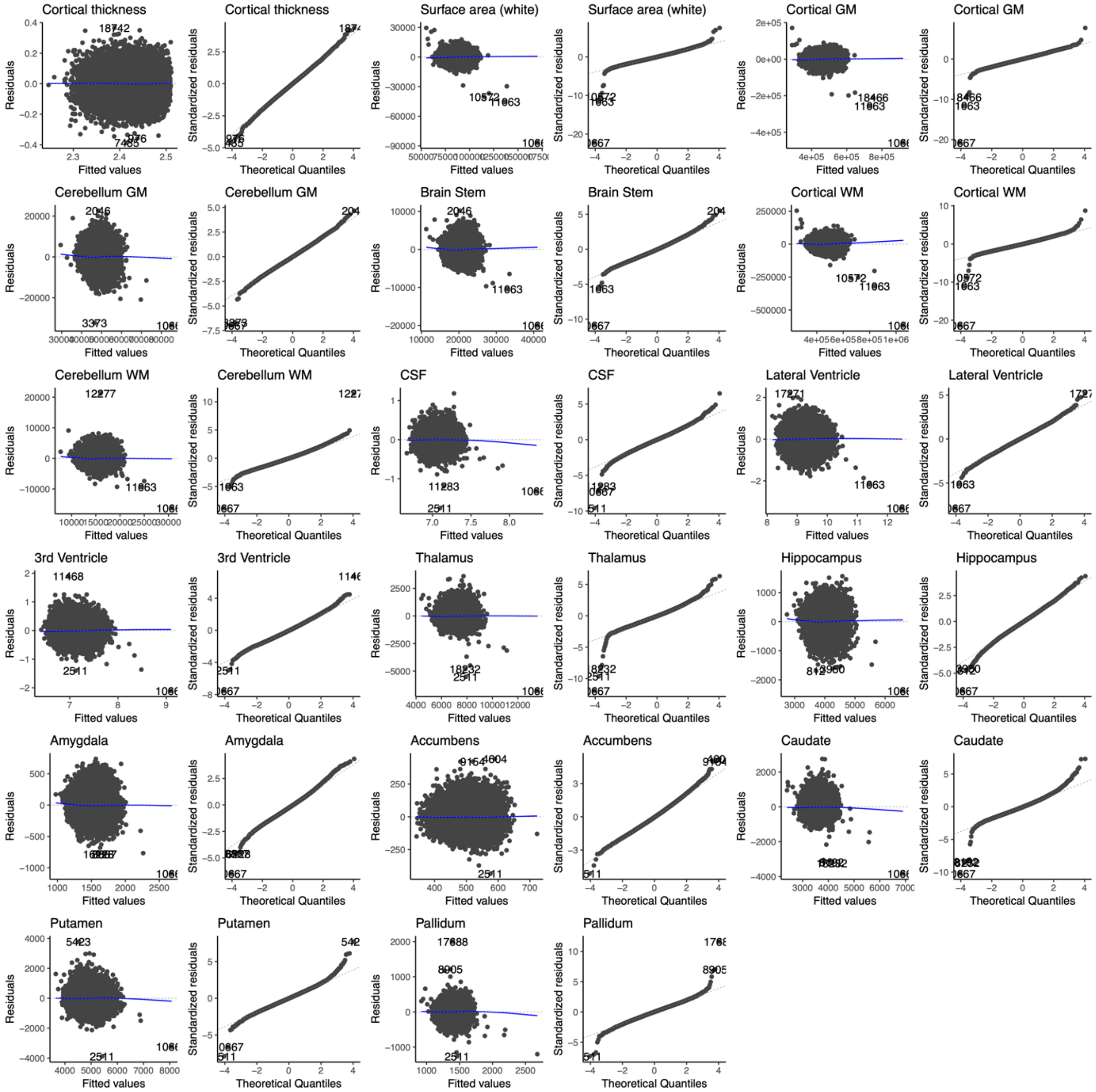


*Notes:* Residuals derived from model 2c with waist-hip-ratio (WHR) as dependent variable. *Abbreviations:* CSF - cerebrospinal fluid; GM – gray matter; WM – white matter.

**Figure S17: Residual versus fitted value plots and Q-Q plots for CSF, lateral ventricle, and 3[rd] ventricle with (left) and without (right) log-transformation (full sample; n=19,330).**

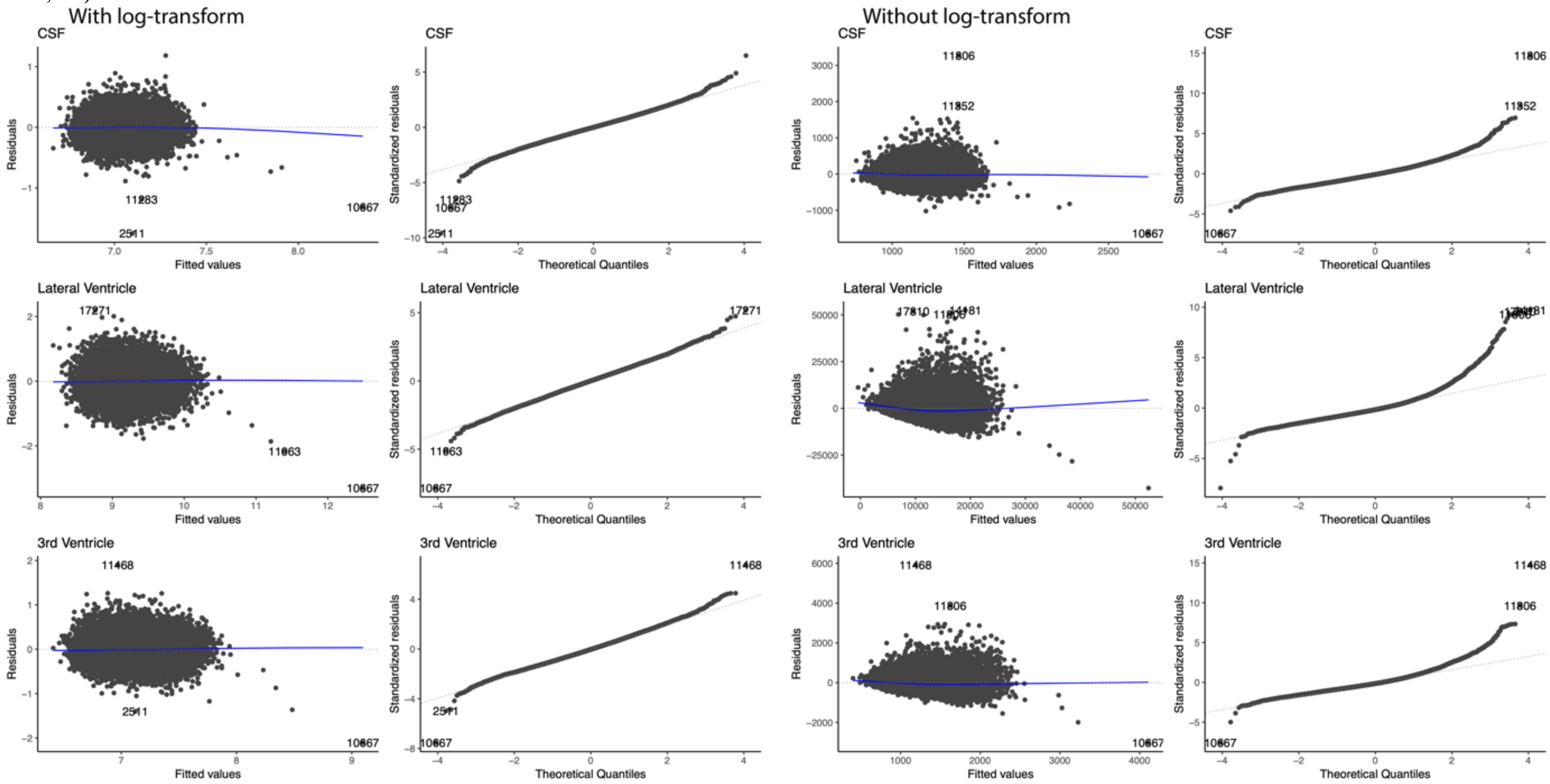


*Notes:* Residuals derived from model 2c with waist-hip-ratio (WHR) as dependent variable. *Abbreviations:* CSF - cerebrospinal fluid.

**Figure S18: Residual versus fitted value plots and Q-Q plots for all body composition models after log-transformation of CSF, lateral ventricle, and 3rd ventricle (body MRI subsample; n=2703).**

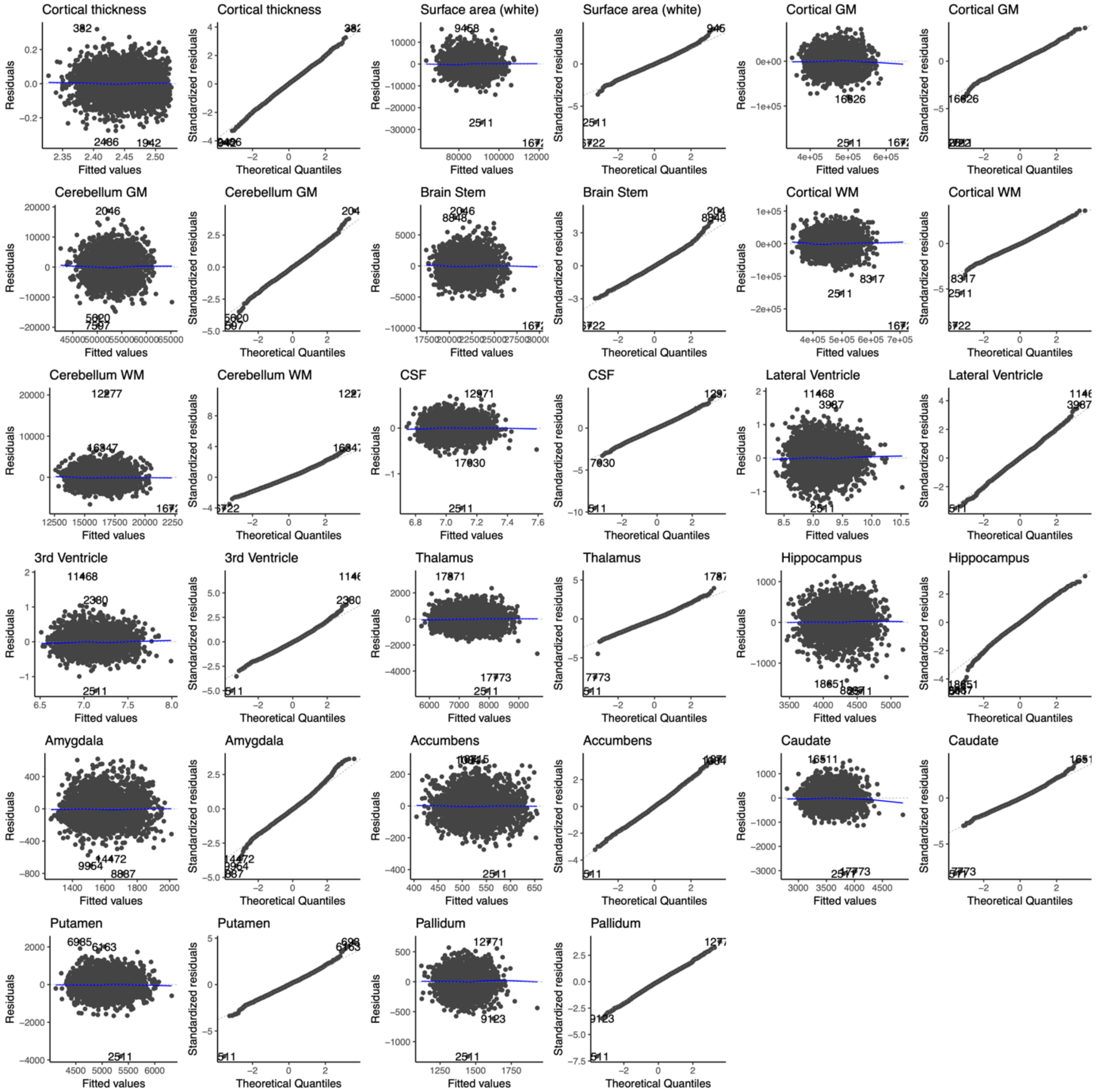


*Notes:* Residuals derived from model 2c with waist-hip-ratio (WHR) as dependent variable. *Abbreviations:* CSF - cerebrospinal fluid; GM – gray matter; WM – white matter.

**Figure S19: Evaluation of multiple linear regression model residuals for normality using residual versus fitted value plots and Q-Q plots for CSF, lateral ventricle, and 3rd ventricle with (left) and without (right) log-transformation (body MRI subsample; n=2703).**

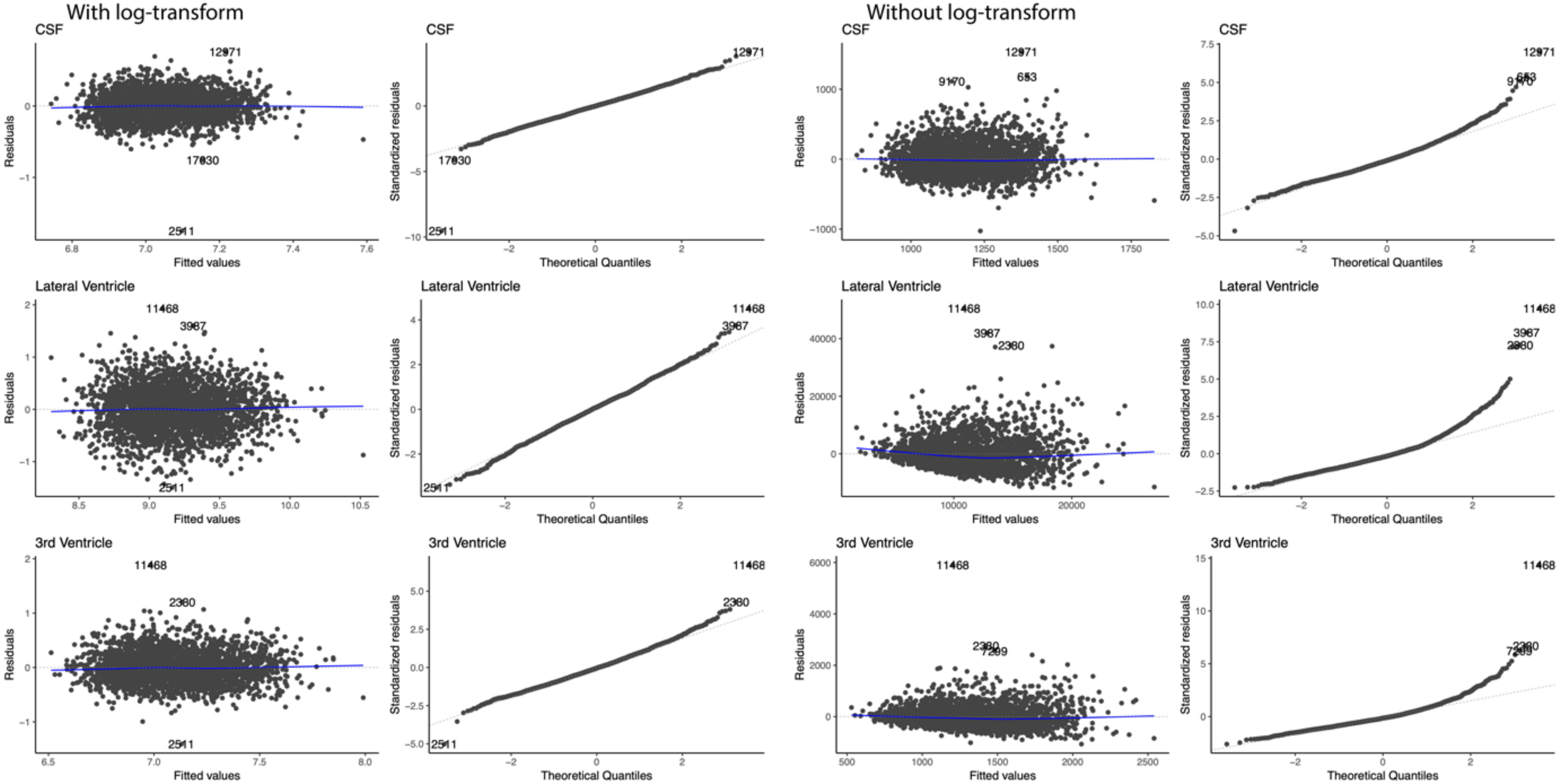


*Notes:* Residuals derived from model 2c with waist-hip-ratio (WHR) as dependent variable. *Abbreviations:* CSF - cerebrospinal fluid; GM – gray matter; WM – white matter.

**Figure S20-a: Linear body-brain associations in healthy across models 2a/b/c (n=19,330).**

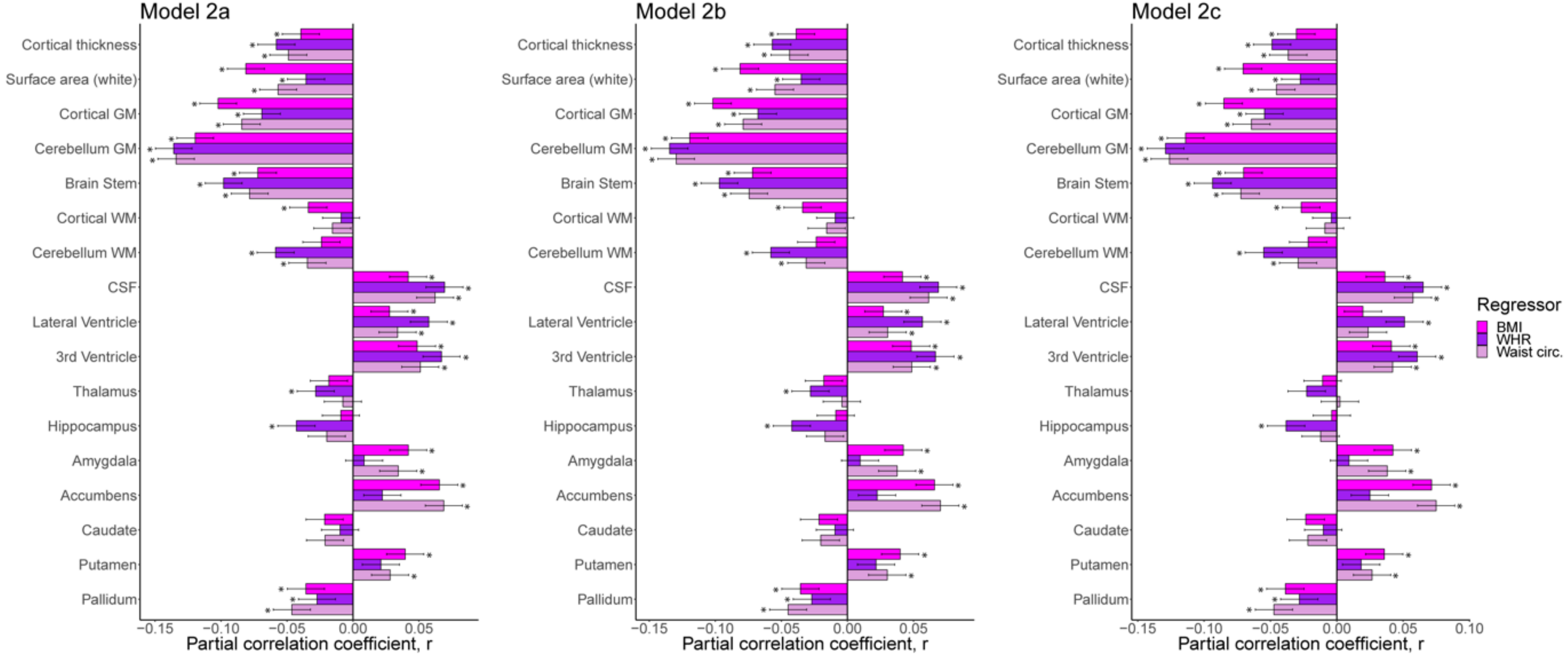


*Notes* Results from model 2a/b/c that investigates body-brain connections through the inclusion of linear and quadratic (only model 2b/c) body composition terms. Figure displays the association between linear body composition term and brain structure. The * indicate significance. Dependent variables CSF, lateral/3$^{rd}$ ventricle were log-transformed. *Abbreviations:* BMI – body mass index; circ. – circumference; GM – gray matter; WHR – waist-to-hip ratio; WM – white matter.

**Figure S20-b: Quadratic body-brain associations in healthy across models 2b/c (n=19,330).**

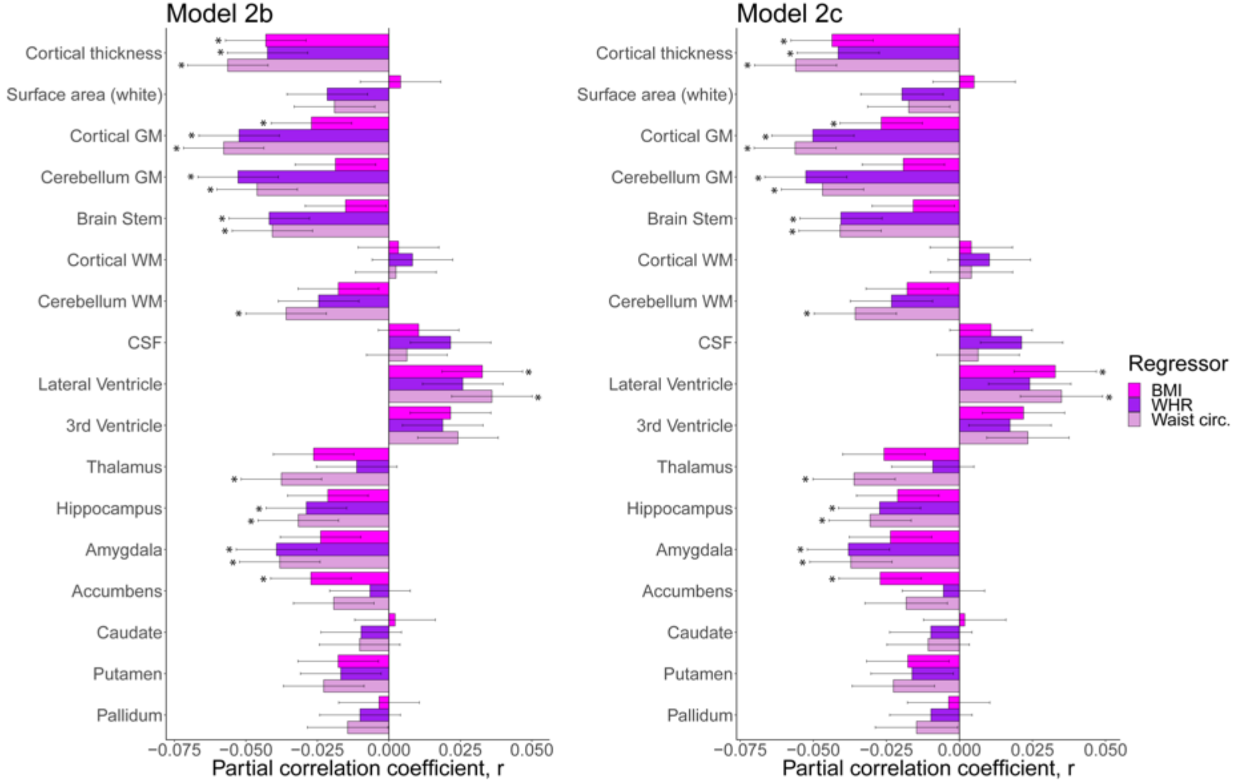


*Notes:* Results from model 2b/c that investigates body-brain connections through the inclusion of linear and quadratic body composition terms. Figure displays the association between quadratic body composition term and brain structure. The * indicate significance. Dependent variables CSF, lateral/3$^{rd}$ ventricle were log-transformed. *Abbreviations:* BMI – body mass index; circ. – circumference; GM – gray matter; WHR – waist-to-hip ratio; WM – white matter.

**Figure S21-a: Linear body-brain associations in healthy across models 2a/b/c (n=2703) for anthropometric measures.**

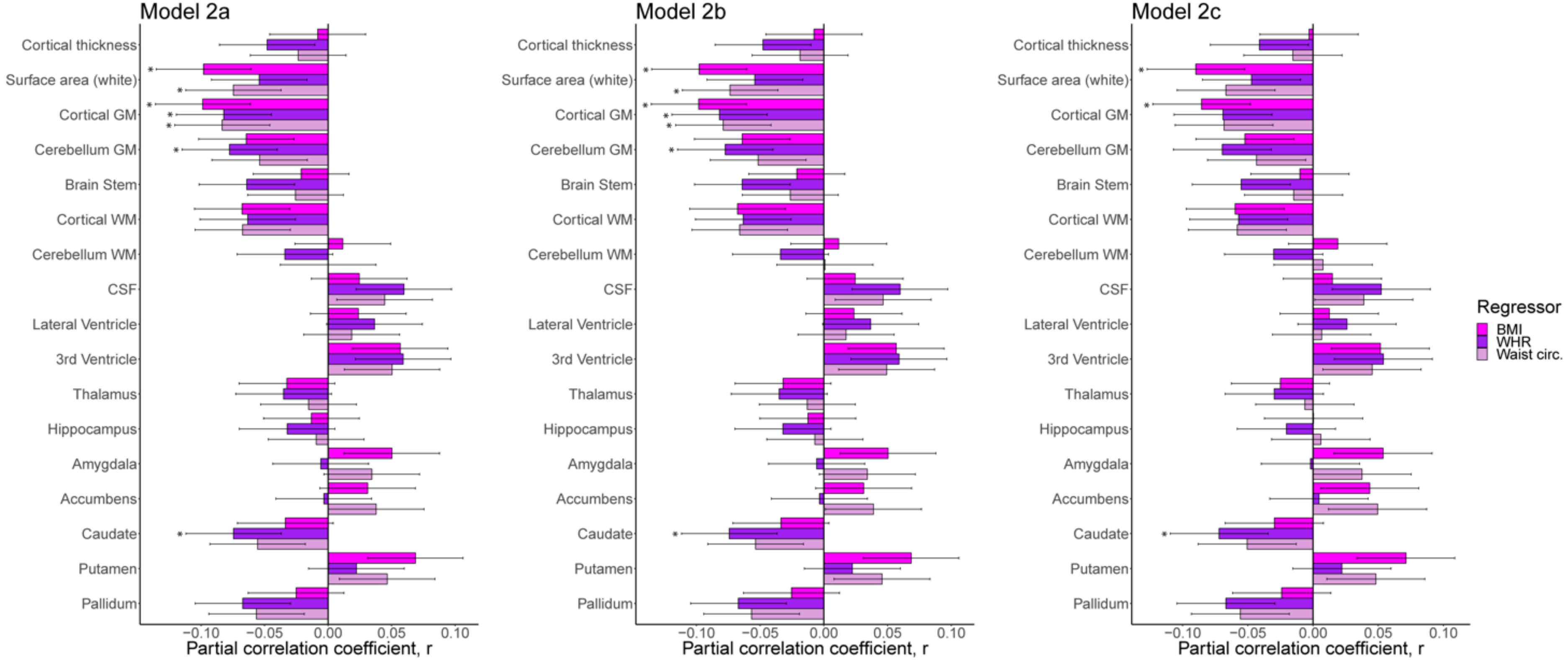


*Notes:* Results from model 2a/b/c that investigates body-brain connections through the inclusion of linear and quadratic (only model 2b/c) body composition terms. Figure displays the association between linear body composition term and brain structure. *Abbreviations:* BMI – body mass index; circ. – circumference; GM- gray matter; WHR – waist-to-hip ratio; WM – white matter.

**Figure S21-b: Quadratic body-brain associations in healthy across models 2b/c (n=2703) for anthropometric measures.**

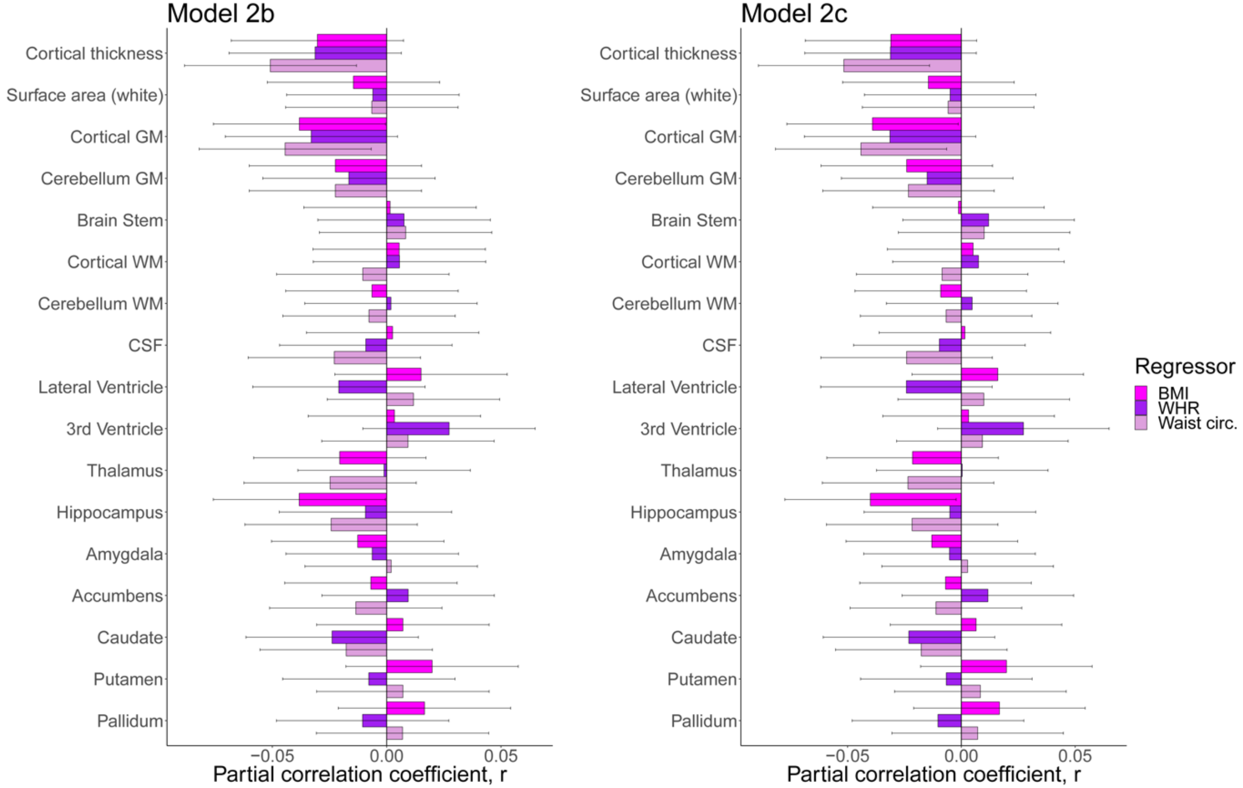


*Notes:* Results from model 2b/c that investigates body-brain connections through the inclusion of linear and quadratic body composition terms. Figure displays the association between quadratic body composition term and brain structure. *Abbreviations:* BMI – body mass index; circ. – circumference; GM- gray matter; WHR – waist-to-hip ratio; WM – white matter.

**Figure S22-a: Linear body-brain associations in healthy across models 2a/b/c (n=2703) for body MRI measures.**

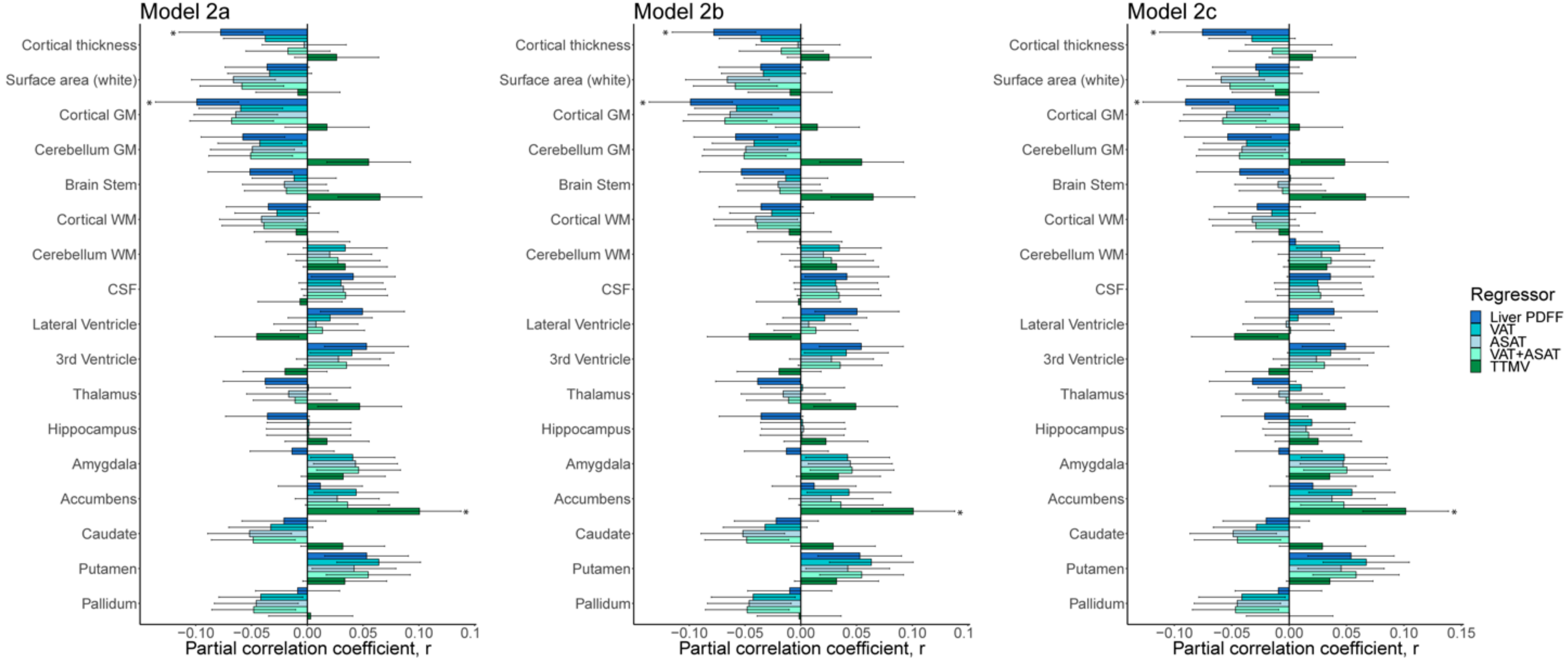


*Notes:* Results from model 2a/b/c that investigates body-brain connections through the inclusion of linear and quadratic (only model 2b/c) body composition terms. Figure displays the association between linear body composition term and brain structure. *Abbreviations:* ASAT – abdominal subcutaneous adipose tissue; GM- gray matter; PDFF – proton density fat fraction; TTMV – total thigh muscle volume; VAT – visceral adipose tissue; WM – white matter.

**Figure S22-b: Quadratic body-brain associations in healthy across models 2b/c (n=2703) for body MRI measures.**

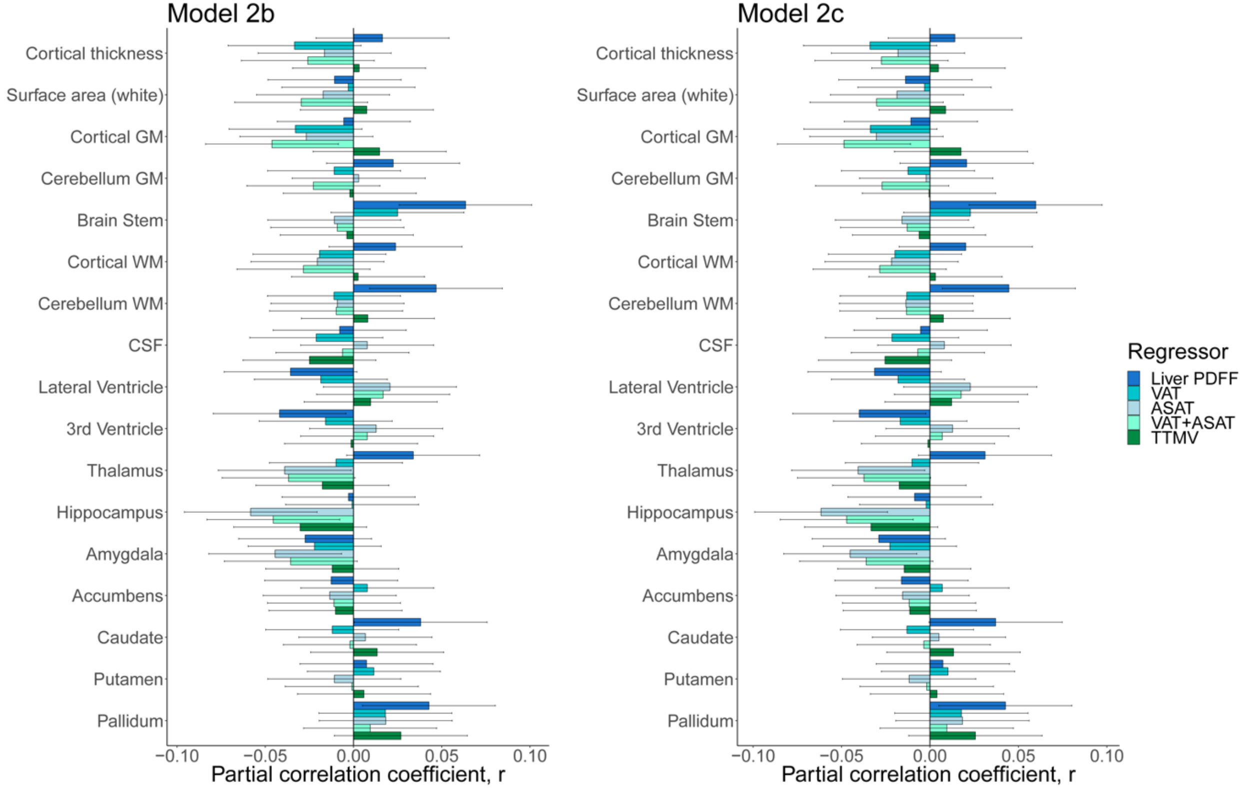


*Notes:* Results from model 2b/c that investigates body-brain connections through the inclusion of linear and quadratic body composition terms. Figure displays the association between quadratic body composition term and brain structure. *Abbreviations:* ASAT – abdominal subcutaneous adipose tissue; GM- gray matter; PDFF – proton density fat fraction; TTMV – total thigh muscle volume; VAT – visceral adipose tissue; WM – white matter.

### Supplemental Tables

**Table S1: Demographics for the body MRI subsample (n=2703).**

| | **Men n=1207** | **Women n=1496** | **$\chi^2$-test/t-test/ Wilcoxon rank-sum test** | **p-value** |
|---|---|---|---|---|
| Age (year) | 61.4±7.4 | 60.5±7.4 | 3 | **0.0024** |
| European ancestry[2] | 1157 (95.9) | 1455 (97.3) | 3.6 | 0.0572 |
| Smoker[2] | 58 (4.8) | 57 (3.8) | 1.4 | 0.2386 |
| Alcohol drinker[2] | 1141 (94.5) | 1400 (93.6) | 0.9 | 0.3412 |
| Height (cm) | 175.8±6.3 | 162.8±6.3 | 53.1 | **0** |
| Weight (kg)[3] | 82.3±11.6 | 68.7±12.4 | 29.3 | **7.8e−164** |
| BMI[3] | 26.6±3.5 | 25.9±4.5 | 4.7 | **3.3e−06** |
| Waist circumference (cm) [3] | 91.8±8.8 | 81±10.7 | 28.9 | **6.3e−160** |
| Hip circumference (cm) [3] | 100.8±6.3 | 100.7±9.3 | 0.6 | 0.5427 |
| WHR[3] | 0.9±0.1 | 0.8±0.1 | 48.1 | **0** |
| Liver PDFF[4] | 4.2±4.6 | 3.3±4.4 | 0.531 | **4.0e−22** |
| VAT[4] | 4.6±2.1 | 2.5±1.4 | 2.006 | **6.3e−157** |
| ASAT[4] | 5.6±2.2 | 7.8±3.3 | −2.0368 | **2.4e−84** |
| VAT+ASAT[3] | 10.1±3.9 | 10.3±4.5 | −0.9 | 0.3474 |
| TTMV[3] | 12.5±1.7 | 8.3±1.2 | 70.5 | 0 |
| Diabetes[2] | 19 (1.6) | 13 (0.9) | 2.3 | 0.132 |
| Hypercholesterolemia[2] | 128 (10.6) | 83 (5.5) | 23 | **1.6e−06** |
| Hypertension[2] | 291 (24.1) | 265 (17.7) | 16.3 | **5.3e−05** |

*Notes*: Report mean ± standard deviation for continuous variables, number of participants (percentage) for categorical variables. P-values<0.05 considered significant. *Abbreviations:* ASAT – abdominal subcutaneous adipose tissue; BMI – body mass index; PDFF – proton density fat fraction; VAT – visceral adipose tissue; WHR – waist-to-hip ratio.

[1] Two-sample t-test

[2] $\chi^2$-test used.

[3] Welch two sample t-test.

[4] Wilcoxon rank-sum test.

**Tables S2-S15 are presented sheet-wise in a separate supplemental excel document.**

The included tables are:

Table S2: Sample description – Age- and sex-related associations on body composition (full sample; n=19,330).

Table S3: Sample description – Age- and sex-related associations on body composition (body MRI subsample; n=2703).

Table S4: Sample description – Age- and sex-related associations on brain structure (full sample; n=19,330).

Table S5: Body-brain associations for BMI as covariate of interests (full sample; n=19,330)

Table S6: Body-brain associations for WHR as covariate of interests (full sample; n=19,330)

Table S7: Body-brain associations for waist circumference as covariate of interests (full sample; n=19,330)

Table S8: Body-brain associations for BMI as covariate of interests (body MRI subsample; n=2703)

Table S9: Body-brain associations for WHR as covariate of interests (body MRI subsample; n=2703)

Table S10: Body-brain associations for waist circumference as covariate of interests (body MRI subsample; n=2703)

Table S11: Body-brain associations for liver proton density fat fraction (PDFF) as covariate of interests (body MRI subsample; n=2703)

Table S12: Body-brain associations for visceral adipose tissue (VAT) as covariate of interests (body MRI subsample; n=2703)

Table S13: Body-brain associations for abdominal subcutaneous adipose tissue (ASAT) as covariate of interests (body MRI subsample; n=2703)

Table S14: Body-brain associations for total abdominal adipose tissue (VAT+ASAT) as covariate of interests (body MRI subsample; n=2703)

Table S15: Body-brain associations for total thigh muscle volume (TTMV) as covariate of interests (body MRI subsample; n=2703)

### Supplemental Notes

#### Note S1: Exclusion criteria for the study

All participants with cancer diagnosis were excluded (http://biobank.ctsu.ox.ac.uk/showcase/; UK-biobank data-field ID 20001). We also excluded participants that were diagnosed with selected traumas, neurological, psychiatric, substance abuse, cardiovascular, liver, or severe infectious conditions. *Table SN1* yields an overview of the excluded non-cancer diagnosis (UK-biobank data-field 20002). We based the exclusion criteria on self-reported diagnosis.

**Table SN1: Excluded non-cancer diagnosis.**

| Code | Meaning | Code | Meaning | Code | Meaning |
|---|---|---|---|---|---|
| **1066** | heart/cardiac problem | **1592** | aortic dissection | **1261** | multiple sclerosis (MS) |
| **1075** | heart attack/myocardial infarction | **1493** | other venous/lymphatic disease | **1262** | parkinsons disease |
| **1076** | heart failure/pulmonary odema | **1494** | varicose veins | **1263** | dementia/alzheimers/cognitive impairment |
| **1077** | heart arrhythmia | **1495** | lymphoedema | **1397** | other demyelinating disease (not MS) |
| **1471** | atrial fibrillation | **1593** | varicose ulcer | **1264** | Epilepsy |
| **1483** | atrial flutter | **1136** | liver/biliary/pancreas problem | **1265** | migraine |
| **1484** | wolff parkinson white/wpw syndrome | **1155** | hepatitis | **1433** | cerebral palsy |
| **1485** | irregular heart beat | **1156** | infective/viral hepatitis | **1434** | other neurological problem |
| **1486** | sick sinus syndrome | **1157** | non-infective hepatitis | **1436** | headaches (not migraine) |
| **1487** | svt/supraventricular tachycardia | **1578** | hepatitis a | **1437** | myasthenia gravis |
| **1488** | mitral valve prolapse | **1579** | hepatitis b | **1525** | benign/essential tremor |
| **1489** | mitral stenosis | **1580** | hepatitis c | **1526** | polio/poliomyelitis |
| **1584** | mitral valve disease | **1581** | hepatitis d | **1659** | meningioma/benign meningeal tumour |
| **1585** | mitral regurgitation/incompetence | **1582** | hepatitis e | **1683** | benign neuroma |
| **1078** | heart valve problem/heart murmur | **1158** | liver failure/cirrhosis | **1240** | neurological injury/trauma |
| **1586** | aortic valve disease | **1506** | primary biliary cirrhosis | **1266** | head injury |
| **1587** | aortic regurgitation/incompetence | **1604** | alcoholic liver disease/alcoholic cirrhosis | **1267** | spinal injury |
| **1490** | aortic stenosis | **1507** | haemochromatosis | **1297** | muscle/soft tissue problem |
| **1079** | cardiomyopathy | **1508** | jaundice (unknown cause) | **1407** | Burns |
| **1588** | hypertrophic cardiomyopathy (hcm/hocm) | **1475** | sclerosing cholangitis | **1626** | fracture skull/head |
| **1589** | pericarditis | **1244** | infection of nervous system | **1630** | fracture neck/cervical fracture |
| **1590** | pericardial effusion | **1245** | brain abscess/intracranial abscess | **1625** | cellulitis |
| **1080** | pericardial problem | **1246** | encephalitis | **1350** | polycystic ovaries/ovarian syndrome |
| **1426** | myocarditis | **1247** | meningitis | **1371** | sarcoidosis |
| **1479** | rheumatic fever | **1248** | spinal abscess | **1439** | hiv/aids |
| **1081** | stroke | **1249** | cranial nerve problem/palsy | **1577** | typhoid fever |
| **1086** | subarachnoid haemorrhage | **1250** | bell's palsy/facial nerve palsy | **1443** | schistosomiasis/bilharzia |
| **1491** | brain haemorrhage | **1523** | trigemminal neuralgia | **1288** | nervous breakdown |
| **1583** | ischaemic stroke | **1251** | spinal cord disorder | **1289** | schizophrenia |
| **1231** | post-natal depression | **1408** | alcohol dependency | **1409** | opioid dependency |
| **1082** | transient ischaemic attack (tia) | **1252** | paraplegia | **1290** | deliberate self-harm/suicide attempt |
| **1083** | subdural haemorrhage/haematoma | **1524** | spina bifida | **1291** | mania/bipolar disorder/manic depression |
| **1425** | cerebral aneurysm | **1254** | peripheral nerve disorder | **1531** | post-natal depression |
| **1067** | peripheral vascular disease | **1256** | acute infective polyneuritis/guillain-barre syndrome | **1469** | post-traumatic stress disorder |
| **1068** | venous thromboembolic disease | **1257** | trapped nerve/compressed nerve | **1470** | anorexia/bulimia/other eating disorder |
| **1087** | leg claudication/intermittent claudication | **1468** | diabetic neuropathy/ulcers | **1615** | Obsessive compulsive disorder (ocd) |
| **1088** | arterial embolism | **1258** | chronic/degenerative neurological problem | **1243** | psychological/psychiatric problem |
| **1093** | pulmonary embolism ± dvt | **1259** | motor neurone disease | **1286** | Depression |
| **1094** | deep venous thrombosis (dvt) | **1260** | myasthenia gravis | **1287** | anxiety/panic attacks |
| **1591** | aortic aneurysm rupture | **1255** | peripheral neuropathy | **1288** | nervous breakdown |
| **1410** | other substance abuse/dependency | **1614** | stress | **1616** | insomnia |

**Note S2: Extracted/computed demographic and clinical variables**

For an overview of of the extracted demographic and clinical data-field IDs from the UK-biobank, see *Table SN2*. Additionally, we computed the following ratios: (1) Body-mass-index (BMI) from weight and standing height as: weight in kg/(height in meters)$^2$, and (2) waist-to-hip ratio computed as: waist circumference/hip circumference.

We created binary (yes/no) variables respectively for: diagnosis (data-field 20002.2.*) of diabetes (diabetes, diabetes type 1, diabetes type2), hypertension, and high cholesterol (hypercholesterolemia); current alcohol consumption, and current cigarette smoking (yes: current, no: previous/never). For ethnicity, we created a binary variable for self-identified European/non-European ancestry based on the MRI time point when available, and complemented incomplete data with baseline information since ethnicity does not change with time (although knowledge/perception of ethnic background can change).

**Table SN2: Extracted demographic and clinical variables with data-field ID.**

| **Data-field ID** | **Field Description** |
|---|---|
| 31.0.0 | Sex |
| 54.2.0 | Assessment center |
| 21003.2.0 | Age |
| 50.2.0 | Standing height |
| 21002.2.0 | Weight |
| 48.2.0 | Waist circumference |
| 49.2.0 | Hip circumference |
| 21000.*.0[1] | Ethnicity |
| 20001.2.*[2] | Cancer diagnosis (self-reported) |
| 20002.2.*[3] | Non-cancer diagnosis (self-reported) |
| 20117.2.0 | Alcohol drinking status |
| 20116.2.0 | Cigarette smoking status |

[1] Baseline and imaging timepoint extracted.
[2] All sub-items (*), from 0 to 5 extracted, imaging timepoint used when available.
[3] All sub-items (*), from 0 to 32 extracted.

**Note S3: MRI acquisition**

Brain MRI was available from three sites (Cheadle, Reading, and Newcastle), and body and liver MRI from one site (Cheadle). Similar scanners/protocols were used across sites.[1,2] Briefly, a single sagittal T1-weighted brain MRI was acquired on a 3T Siemens Skyra scanner equipped with a 32-channel head coil using a 3D MPRAGE sequence with pre-scan normalization.[1,2] Body and liver MRI was acquired on a 1·5T Siemens MAGNETOM Aera scanner using a dual-echo Dixon Vibe protocol[3] and a single transverse breath-hold multi-echo spoiled-gradient-echo acquisition without contrast agent,[4] respectively.

**Note S4: Body MRI processing details**

For the body MRI data, we acquired the processed data from the UK-biobank. The data was processed for abdominal fat and thigh muscle volume by AMRA (Linköping, Sweden; https://www.amramedical.com),[3] and liver proton density fat fraction (PDFF) by Perspectum Diagnostics (Oxford, UK; https://perspectum-diagnostics.com).[4]

The body MRI processing by AMRA include intensity inhomogeneity correction, non-rigid registration of atlases to acquired image volumes, quantification of fat and muscle composition using a voting scheme, and visual inspection for segmentation accuracy and manual adjustment[3] (for technical details, see[5–7]). Adipose tissue within the abdominal cavity was defined as visceral adipose tissue (VAT), adipose tissue between the top of the femoral head and the top of T9 was defined as abdominal subcutaneous adipose tissue (ASAT), and lean thigh muscle volume included the gluteus, iliacus, adductors, hamstrings, quadriceps femoris and sartorius.[3] AMRA implements manual quality control of the image/segmentation quality.

The liver MRI was processed by Perspectum Diagnostics using the LiverMultiScan$^{TM}$ Discovery software that utilize the $2^{nd}$, $4^{th}$, and $6^{th}$ of 10 MRI echoes from three-point DIXON technique to construct liver proton density fat fractions (PDFF). Three circular regions of interests (ROIs) with 15 mm diameter were selected from the PDFF map. The reported PDFF is the mean of all pixels within the three ROIs. Perspectum Diagnostics implements manual quality control of the image/segmentation quality.[4] The PDFF is computed as *fat/(water+fat)* (http://biobank.ctsu.ox.ac.uk/showcase/).

We extracted volumetric body MRI variables from the UK-biobank repository (*Table SN3*). For further information about the UK-biobank body MRI, see the documentation at the UK-biobank showcase (http://biobank.ctsu.ox.ac.uk/showcase/).

**Table SN3: Extracted body MRI variables with data-field ID**

| Data-field ID | Field Description |
|---|---|
| 12224.2.0 | Indications whether abdominal MRI has been completed. |
| 22402.2.0[1] | Liver fat percentage |
| 22407.2.0 | Visceral adipose tissue (VAT) volume |
| 22408.2.0 | Abdominal subcutaneous adipose tissue (ASAT) volume |
| 22409.2.0 | Total thigh muscle volume (TTMV; sum of all thigh lean muscle volumes) |
| 22410.2.0 | Total trunk fat (VAT+ASAT) |
| 22414.2.*[2] | Image quality indicator |

[1] Field yields liver proton density fat fraction (PDFF).
[2]All sub-items extracted (0 and 1). Variable used for quality control of body MRI measures.

**Note S5: Brain MRI Quality control**

For the T1 weighted brain MRI data, we applied an automated quality control based on the FreeSurfer[8] Euler number.[9,10] Higher negative Euler number imply worse image quality. For each hemisphere, participants were iteratively excluded if the Euler number exceeded three standard deviations (SD) from the mean in either hemisphere (one-sided). We iterated until there were no outliers left, resulting in eight iterations (*Figure S1*).

*Table SN4* gives the number of excluded/included participants together with the average Euler number, while *Figure SN1* shows illustrates the Euler number distribution of included/excluded participants for the left/right hemisphere.

**Table SN4: Average FreeSurfer Euler number of excluded/included participants.**

| | **Included (N=19 330)** | **Excluded (N=2 063)** |
|---|---|---|
| Left hemisphere | -54.1±24.9 | -182.8±90.8 |
| Right hemisphere | -51.7±23.7 | -172.6±87 |

*Notes:* Report mean ± standard deviation

**Figure SN1: Violin plots of included/excluded participants based FreeSurfer Euler numbers.**

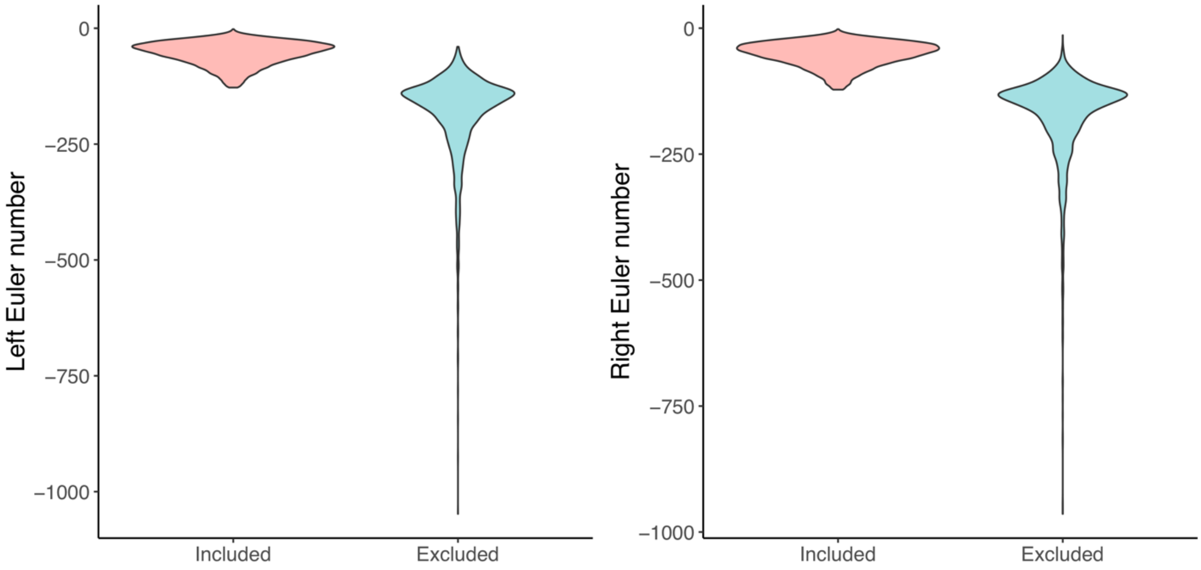


*Notes:* Plots of included/excluded participants are not adjusted for sample size.

**Note S6: Linear regression models using the *lm* in r**

Below we present the linear regression models as implemented in R (version 3·5·2; https://www.r-project.org) using the *lm* for the sample description and main analyses.

*Sample description: Body composition*

**Model 1a: lm(**log(body[†]) ~ poly(Age,2)+Sex**)**

**Model 1b: lm(**log(body[†]) ~ poly(Age,2)×Sex**)**

**Model 1c: lm(**log(body[†]) ~ poly(Age,2)×Sex + Ethnicity + Diabetic + Hypercholesteremia + Hypertension + Smoking + Alcohol**)**

*Sample description: Brain structure*

**Model 1a: lm(**brain[‡] ~ poly(Age,2)+Sex + ICV[§] + Euler number + Assessment center**)**

**Model 1b: lm(**brain[‡] ~ poly(Age,2)×Sex + ICV[§] + Euler number + Assessment center**)**

*Brain structure and body composition measures*

***Model 2a: lm(****brain[‡] ~ body[†] + poly(Age,2)×Sex + ICV[§] + Euler number + Assessment center[**]**)***

***Model 2b: lm(****brain[‡] ~ poly(body[†],2) + poly(Age,2)×Sex + ICV[§] + Euler number + Assessment center[**]**)***

***Model 2c: lm(****brain[‡] ~ poly(body[†],2) + poly(Age,2)×Sex + ICV[§] + Ethnicity + Diabetic + Hypercholesteremia+ Hypertension + Smoking + Alcohol + Euler number + Assessment center[**]**)***

[†] Body composition measure.
[‡] Measured brain structure. We *log*-transformed CSF, Lateral and 3rd ventricle.
[§] We did not adjust the mean cortical thickness for ICV (intracranial volume).
[**] Not included for body MRI measures (single site).

**Note S7: Sample description analyses of body composition and brain structure.**

*Sample description: Body composition*

Analyses including the full sample (*Figure SN2*; *Table S2*) revealed, as expected, higher BMI, WHR, and waist circumference in men compared to women, and that age was negatively associated with BMI and positively with WHR and waist circumference. In the body MRI subsample (*Figure SN3*; *Table S3*), men showed higher liver PDFF, VAT, and TTMV, and lower ASAT compared to women. Age was positively associated with liver PDFF and VAT (significant age-by-sex interactions indicate attenuation in men), and negatively with TTMV. These effects were similar across models 1a/b/c.

*Sample description: Brain structure*

Analyses including the full sample revealed significant age- and sex-associations across most included brain structures. As expected, global cortical and cerebellum measures and subcortical structures showed age-related decreases with significant quadratic terms, indicating increasing age-related associations at higher ages, while CSF, lateral, and third ventricles showed similar increases with age. Men generally showed larger brain volumes and steeper age-related decreases, except for attenuation of age-related ventricular expansion. These effects were similar across models 1a/b (*Figure SN4*; *Table S4*).

**Figure SN2: Age and sex in relation to body composition measures (n=19,330).**

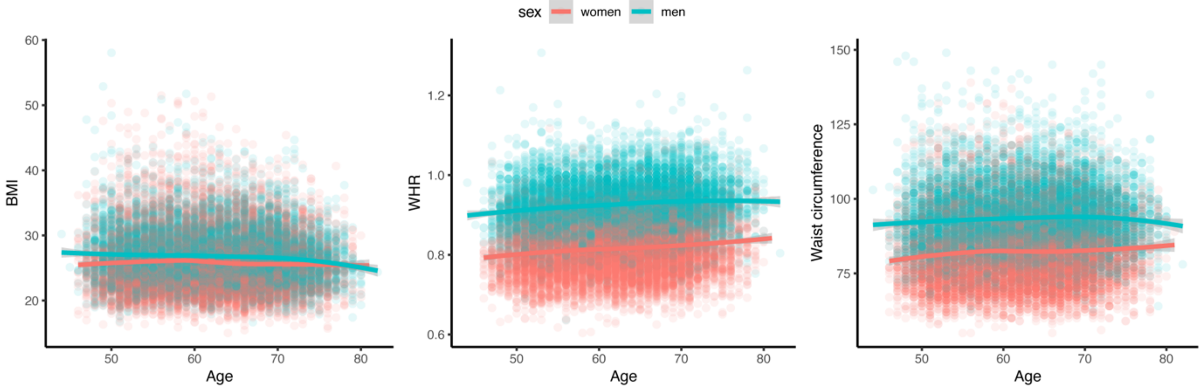


*Notes:* Unadjusted LOESS scatter plots. *Abbreviations:* BMI – Body mass index; WHR – waist-hip-ratio.

**Figure SN3: Age and sex in relation to body composition measures (n=2703).**

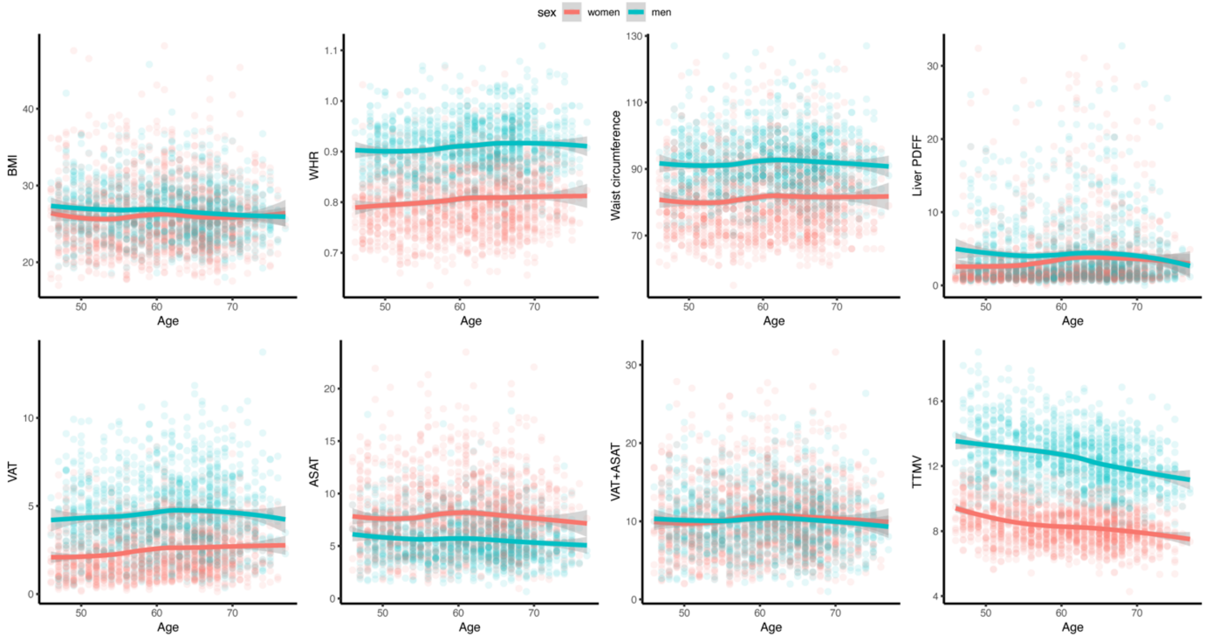


*Notes:* Unadjusted LOESS scatter plots. *Abbreviations:* ASAT – abdominal subcutaneous adipose tissue; BMI – Body mass index; PDFF – proton density fat fraction; TTMV – total thigh muscle volume; VAT – visceral adipose tissue; VAT+ASAT – total abdominal adipose tissue; WHR – waist-hip-ratio.

**Figure SN4: Age and sex in relation to brain structure in generally healthy participants (n=19,330).**

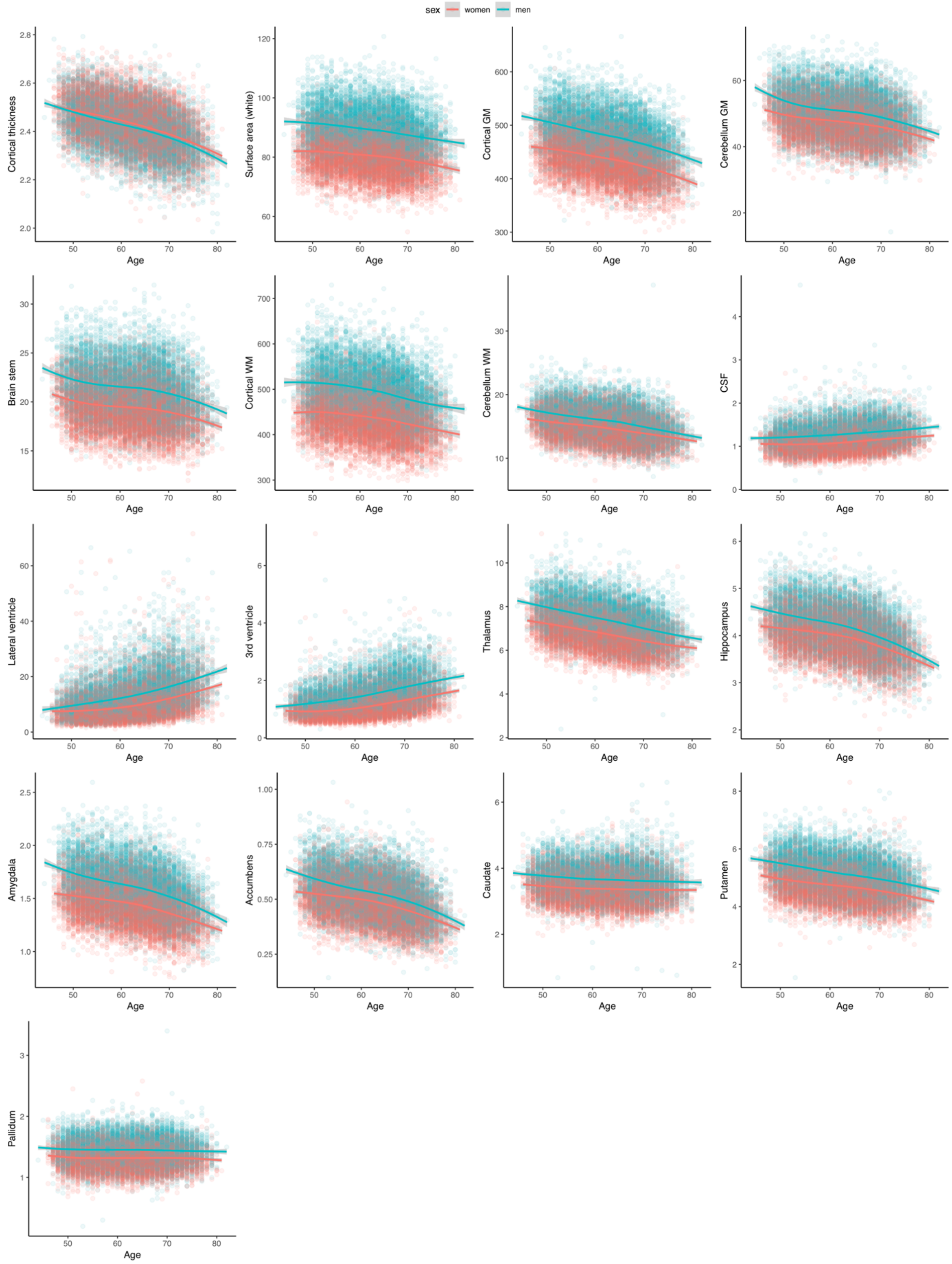


*Notes*: Unadjusted LOESS scatter plots. All structures given in ml (except surface area given in $m^2$). *Abbreviations:* CSF – cerebrospinal fluid; GM – gray matter; ICV – intracranial volume; WM – white matter.
